# Crosstalk and Ultrasensitivity in Protein Degradation Pathways

**DOI:** 10.1101/594085

**Authors:** Abhishek Mallela, Maulik K. Nariya, Eric J. Deeds

## Abstract

Protein turnover is vital to protein homeostasis within the cell. Many proteins are degraded efficiently only after they have been post-translationally “tagged” with a polyubiquitin chain. Ubiquitylation is a form of Post-Translational Modification (PTM): addition of a ubiquitin to the chain is catalyzed by E3 ligases, and removal of ubiquitin is catalyzed by a De-UBiquitylating enzyme (DUB). Over three decades ago, Goldbeter and Koshland discovered that reversible PTM cycles function like on-off switches when the substrates are at saturating concentrations. Although this finding has had profound implications for the understanding of switch-like behavior in biochemical networks, the general behavior of PTM cycles subject to synthesis and degradation has not been studied. Using a mathematical modeling approach, we found that simply introducing protein turnover to a standard modification cycle has profound effects, including significantly reducing the switch-like nature of the response. Our findings suggest that many classic results on PTM cycles may not hold *in vivo* where protein turnover is ubiquitous. We also found that proteins sharing an E3 ligase can have closely related changes in their expression levels. These results imply that it may be difficult to interpret experimental results obtained from either overexpressing or knocking down protein levels, since changes in protein expression can be coupled via E3 ligase crosstalk. Understanding crosstalk and competition for E3 ligases will be key to ultimately developing a global picture of protein homeostasis.

## Introduction

All proteins undergo some form of turnover. For instance, proteins can become damaged via deamidation or some other process and must be degraded in order to prevent unfolding and aggregation. Turnover is also important in signaling and the regulation of protein function. A classic example is the degradation of *IκB* proteins, which bind the NF-*κB* protein complex and sequester it in the cytoplasm. During response to different stimuli, *IκB*s are phosphorylated by *IκB* kinases, ubiquitylated, and tagged for degradation, which allows NF-*κB* to translocate to the nucleus (1). In the cell, synthesis and degradation (i.e. protein turnover) act in concert to maintain an appropriate concentration of active protein (i.e. protein homeostasis). Given the centrality of protein turnover to all cellular processes, it is not surprising that dysregulation of protein homeostasis has been implicated in a vast array of neurodegenerative diseases and cancers (2, 3). In eukaryotes, degradation is often achieved through the ubiquitin-proteasome system, where proteins are tagged with polyubiquitin chains that are recognized by the proteasome, ultimately leading to protein degradation (4). Polyubiquitylation represents a form of post-translational modification (PTM) cycle where ubiquitin subunits are covalently linked to substrates by E3 ligases and removed by deubiquitylating (DUB) enzymes (5).

Over 35 years ago, Goldbeter and Koshland studied the general properties of a PTM cycle comprised of a modifying and demodifying enzyme. They found that reversible cycles of protein modification, such as a kinase enzyme adding a phosphoryl group and a phosphatase enzyme removing it, work like on-off switches when the enzymes are saturated (6). This phenomenon, known as “0^th^-order ultrasensitivity”, has had profound implications for understanding how biochemical networks can exhibit switch-like behavior. Despite decades of progress in understanding 0^th^-order ultrasensitivity and other aspects of PTM function (7–12), to there have been few attempts to systematically characterize the general behavior of PTM cycles that drive protein degradation.

The only exception to this has been the study of ubiquitylation in the context of cell cycle oscillations and bistability (13, 14). While these studies have provided key insights about cell cycle control, they have not investigated how ubiquitylation levels control the steady-state expression levels of proteins not involved in the cell cycle. It has also been shown that adding protein synthesis and degradation to models of gene expression and cell signaling can have dramatic effects on system dynamics, but the detailed impact of turnover on PTM cycles remains unclear (15–17).

In addition to a general lack of understanding of the influence of protein homeostasis on PTM cycle behavior, we recently discovered that substrates in such cycles can have coupled steady-state responses if those substrates share modification/demodification enzymes. In particular, if one substrate is at saturating levels, or if the substrates collectively saturate the enzymes, then all substrates of that pair will respond in a coupled, switch-like manner (18–20). This implies that modification leading to substrate degradation (e.g. ubiquitylation by an E3 ligase) could introduce coupling in the concentrations of substrates sharing a ligase. Interestingly, Mather and co-workers have shown that substrate concentrations can be coupled through saturation of the downstream degradation machinery (21, 22). It is currently unclear, though, whether such coupling can arise due to “crosstalk” in the upstream mechanisms that tag proteins for degradation.

In this work, we used a set of mathematical models to show that perturbing a standard PTM cycle by simply adding synthesis and degradation has profound effects on the response of the system. Specifically, we found that the sensitivity of the system to incoming signals and the ultrasensitivity of the response are dramatically muted when the substrate is at saturating concentrations. When the modification in question drives protein degradation at a higher rate, these effects are even more pronounced. Furthermore, more realistic models allowing for long ubiquitin chains exhibit qualitatively similar behavior to the case with a single modification state, but with further decreases in sensitivity and ultrasensitivity. These findings are robust to changes in the specific mechanisms utilized by the E3 and DUB enzymes. Interestingly, we found that distinct modes of enzyme saturation (i.e. increasing substrate production rate vs. decreasing the Michaelis constant of the enzyme) also effect substrate responses differently. This indicates that many classic results on PTM cycles, including the extremely ultrasensitive response they exhibit when the substrates are at saturating concentrations, may not hold *in vivo* where protein turnover is inevitable. We also found that proteins sharing an E3 ligase can indeed have closely related expression profiles. Moreover, the sensitivity protein concentration to changes in E3 activity for any given protein is largely dependent upon the total expression level of the other proteins. This suggests that it may be difficult to interpret experimental results obtained from either overexpressing or reducing protein concentrations, since changes in protein expression can be coupled via E3 ligase crosstalk. Further experimental characterization of E3-ligase/DUB enzyme/substrate relationships will thus be vital to developing a global understanding of protein regulation within the cell.

## Results

### Competition among E3 ligases

As mentioned in the introduction, shared E3 ligases have the potential to induce coupling in substrate responses. It is currently unclear, however, how widespread such “crosstalk” among E3 ligases might be. We searched the E3Net database (23) for statistics of E3-substrate interactions in human cells. For sake of comparison, we also obtained E3-specific statistics from the hUbiquitome database (24). The total number of E3 ligases documented in E3Net is 415 and the total number of their substrates is 873, making the average ‘substrate load’ (substrate-to-ligase ratio) 2.10. Similarly, there are a total of 138 ligases and 279 substrates annotated in the hUbiquitome database, yielding a comparable ratio of 2.02. Thus, on average, most E3 ligases will ubiquitylate around two substrates.

In addition to providing the numbers of ligases and substrates, E3Net also captures information on specific E3-substrate interactions. We found that 54% of the E3 ligases in the database have no substrates listed; however, of the remaining E3 ligases, 52% have at least 2 substrates and 11% have more than 10 substrates (Fig. 1a). Also, the maximum number of substrates for any ligase is 92. Given that the database is incomplete, it is likely that these numbers represent significant underestimates of E3 ligase crosstalk. Regardless, the phenomenon of E3 ligases acting on multiple substrates is likely widespread, and little is presently known about what influence crosstalk might have on the responses of these substrates to changes in E3 ligase activity.

**Fig. 1.**
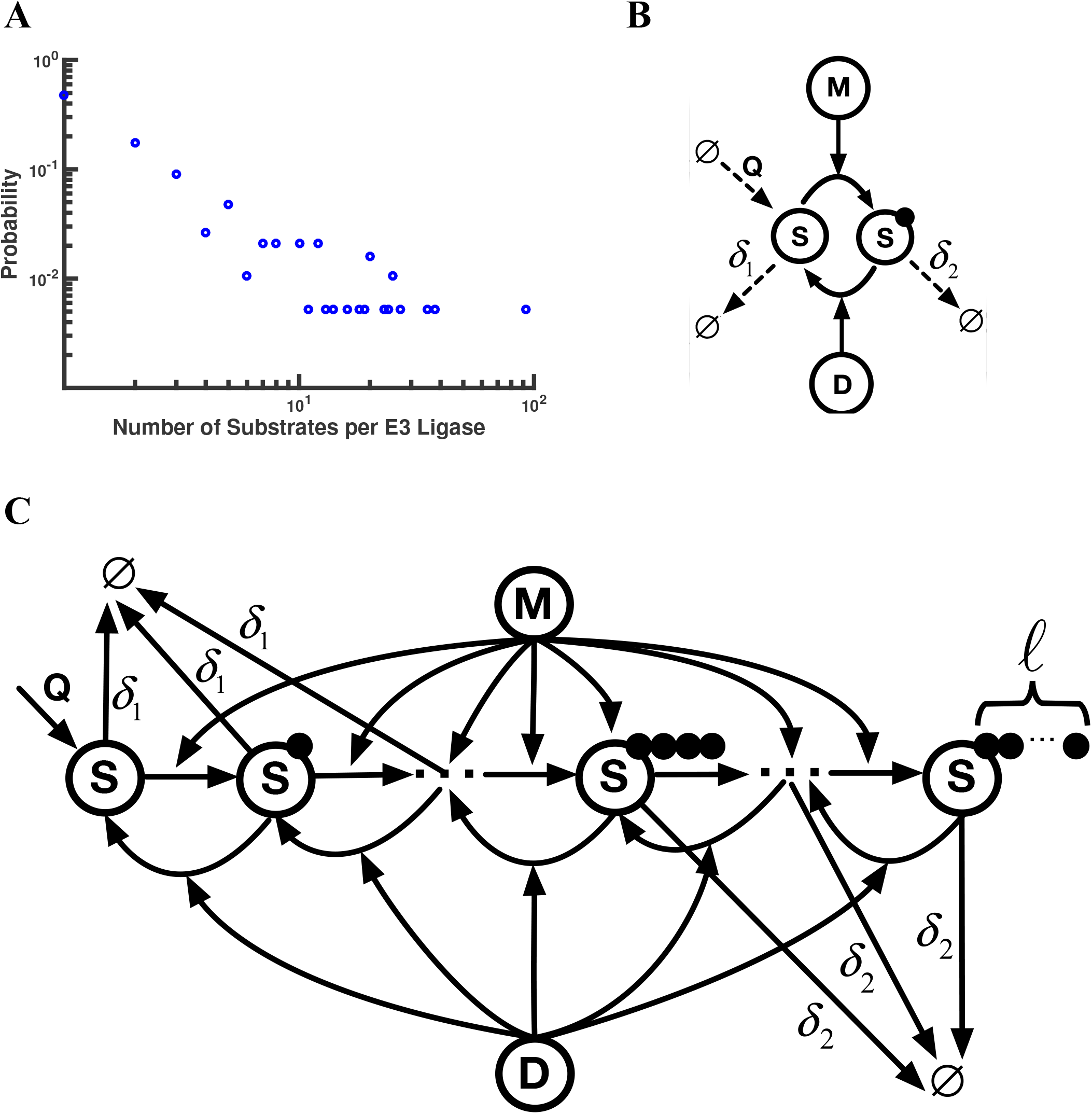
Crosstalk among E3 ligases & schematic diagrams of single-substrate models. **(A)** Probability distribution (on log-log scale) of E3 ligase-substrate specificity as recorded in the E3Net database. The average “substrate load” on a given E3 ligase is 2.1. **(B)** General representation of the canonical Goldbeter-Koshland (GK) loop, with protein turnover included. The case Q = δ_2_ = δ_1_ = 0 corresponds to the GK loop; Q > 0 and δ_2_ = δ_1_ > 0 corresponds to the “Intermediate” model; Q > 0 and δ_2_ > δ_1_ > 0 is represented by the “Full” model. Here “M” denotes modifying enzyme and “D” denotes demodifying enzyme. Modified substrate is indicated by the dark circle. **(C)** Schematic diagram for one substrate with multiple modification states. Shown here is the model corresponding to the Processive E3, Distributive/Sequential DUB case. Each of the first three units is degraded at the rate δ_1_, which is smaller than the rate δ_2_ for the remaining units. The maximum length of the polyubiquitin chain is denoted by *ℓ*.

### Adding synthesis and degradation to a PTM cycle

Even though E3 ligases generally attach long ubiquitin chains to their substrates (4), in order to simplify the problem to an analytically tractable form, we first considered a case with just a single modification state (Fig. 1b vs. 1c). Because ubiquitylation actively effects protein degradation, any investigation of the interplay between substrates competing for a protein and post-translational modifications (PTMs) leading to degradation must account for protein turnover. We thus focused first on studying how synthesis and degradation influence the behavior of the standard Goldbeter-Koshland loop.

The first model, which we call the ‘Intermediate’ model, involves one substrate that can exist in two forms: modified and unmodified, denoted by *S\** and *S* respectively (Fig. 1b). When in the *S* state substrate is degraded at a first-order rate *δ*_1_, and when in the *S\** state it is degraded at a rate *δ*_2_. In the intermediate model, the modification (e.g. ubiquitylation) does not lead to higher rates of degradation, so *δ*_1_ = *δ*_2_. Unmodified substrate is also synthesized at a constant rate *Q*. The enzymatic reaction scheme can be used to obtain a system of ordinary differential equations (ODEs) with the binding, dissociation, and catalysis steps treated explicitly. We have denoted the kinetic rates of complex formation, complex dissociation, and catalysis by *k_x,y_*, where *x* represents the reaction step and *y* represents the enzyme – modifying (M) or demodifying (D) (Supp Info Sec. 1.1). For example, *k_cat,D_* denotes the catalytic rate of the reaction catalyzed by the demodifying enzyme.

Traditional analyses of post-translational modification cycles (i.e. the GK loop) have examined the response of molar fraction of modified protein at steady-state (*S\** ≡ [*S\**]/[*S*]*_T_*, where [*S*]*_T_* = [*S*] + [*S\**] + [*MS*] + [*DS\**]) to changes in the input parameter 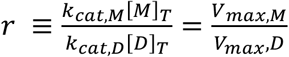 (6, 18). We varied *r* by simply changing [*M*] and numerically integrated the system to extract the steady-state solutions for unmodified and modified substrate ([*S*] and [*S\**]) at each value of *r*. Note that, for the intermediate model, [*S*]*_T_* = *Q*/*δ*_1_, regardless of the values of other parameters (Supp Info Sec. 1.3). Here, we focus on the case where the Michaelis constants of the enzymes are equal (*K_MM_* = *K_M,D_*), leaving analysis of substantially different *K_M_′*s to future work (25).

One key feature of GK loops is their capacity to exhibit 0^th^-order ultrasensitivity, which manifests as a switch-like transition in [*S\**] vs. *r* when the modification and demodification enzymes are saturated (i.e. [*S*]*_T_* ≫ *K_M_*) (6, 9, 18, 25). To explore this phenomenon in the intermediate model, we initially increased *Q* to change saturation levels, since [*S*]*_T_* = *Q*/*δ*_1_. In order to conduct these simulations, we chose a set of reasonable values for the kinetic rate constants in the model (Supp Info Table S1). In particular, *k_cat_*s and *K_M_*s were taken from experimentally observed ranges (Supp Info Figure S1, (26)) and *δ*_1_ was set based on the average observed half-life for proteins in living human cells (27). The value for *δ*_1_ is also very similar to the shortest observed protein half-life in mouse C2C12 cells (20, 28). As can be seen from Fig. 2a, there are dramatic differences between a GK loop and the intermediate model upon saturation. For instance, defining *r*_50_ as the *r*-value when [*S\**] is half-maximal (i.e. [*S\**] = 0.5), we see that the curve of *S\** vs. *r* shifts to the right, indicating a higher *r*_50_. Secondly, the ultrasensitivity (i.e. the effective Hill coefficient *n_eff_*) of the system under the intermediate model is reduced. Since the intermediate model is simply a GK loop with *Q* and *δ*_1_ added (i.e. *δ*_1_ = *δ*_2_), these results indicate that adding synthesis and degradation to a PTM can have a dramatic effect on 0^th^-order ultrasensitivity.

**Fig. 2.**
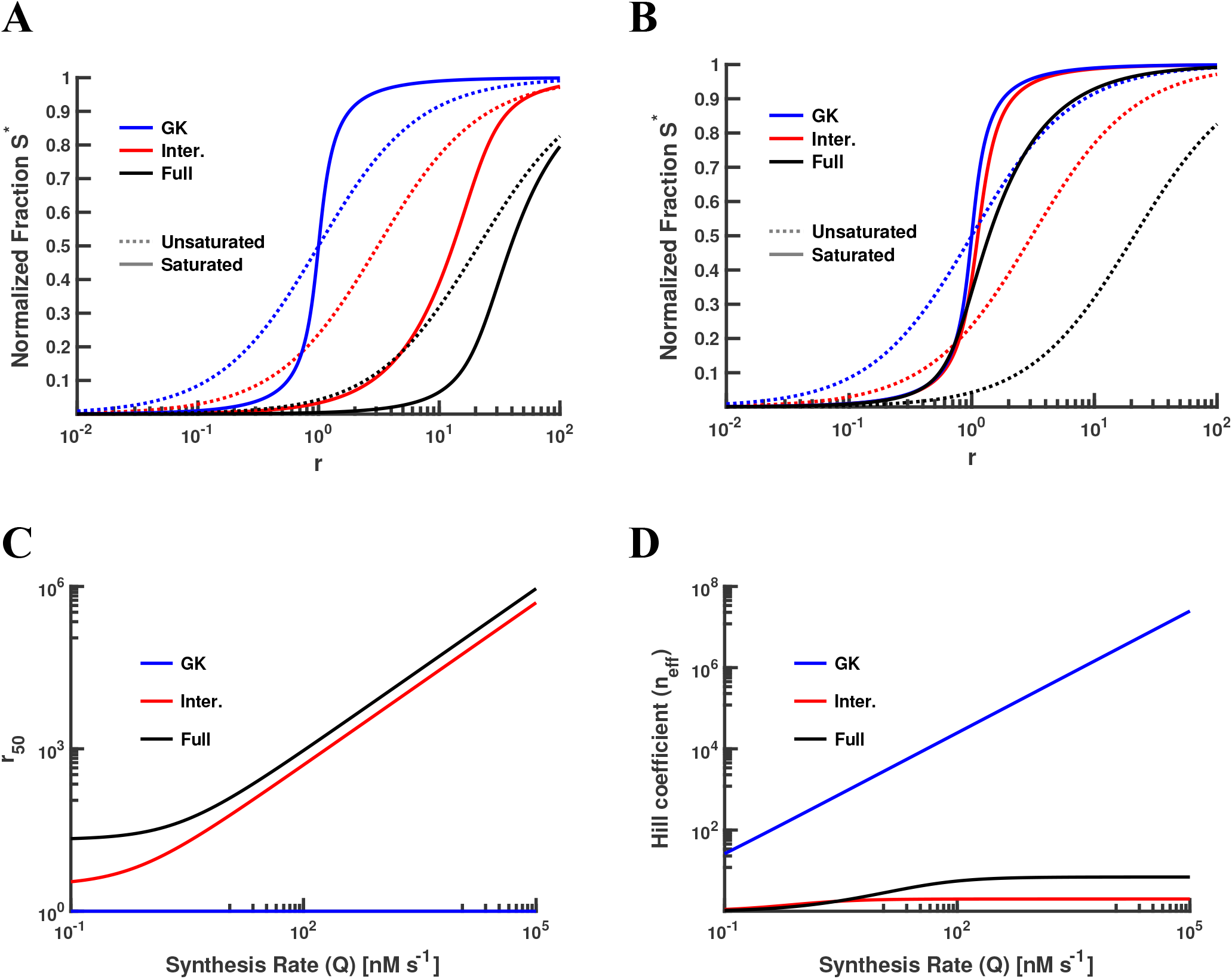
Effects on various single-substrate models of varying Q or K_M_ as the measure of enzyme saturation. **(A)** Modulating the rate of protein synthesis (Q) results in dramatic reduction of both sensitivity of the system to incoming signals and ultrasensitivity of the response, in the regime of saturated enzyme. This is indicated by the rightward shift of the r_50_ and the reduction in the Hill coefficient (n_eff_) from GK to Full. Logarithmic axis used for r. **(B)** Varying the Michaelis-Menten constant (K_M_) results in a smaller reduction of r_S0_ and n_eff_, as compared to varying Q, in the regime of saturated enzyme. **(C)** Increasing Q has no effect on the r_50_ for the GK model; the reduction in sensitivity is highly pronounced for the Intermediate model, and even more so for the Full model. Axes in log scale. **(D)** Increasing Q results in an unbounded increase of n_eff_ for the GK model. However, for systems that incorporate protein turnover (i.e. the Intermediate and Full models), there is a natural limit to the increase in n_eff_ for large enough Q.

While increasing the expression level of *S* (e.g. increasing *Q*) is a natural way to achieve saturation, one can also saturate the enzymes by decreasing *K*_M_, keeping *Q* fixed. In a standard GK loop, varying [*S*]*_T_* and *K_M_* are mathematically equivalent; in the intermediate model, however, the effects of decreasing *K_M_* (with a lower bound of 100 nM for experimentally observed *K_M_*s) are dramatically different from the effects of increasing *Q*. In particular, the change in *r*_50_ and the change in *n_eff_* are negligible (Fig. 2b).

These results can be further understood by treating the system of ODEs analytically at steady-state. We obtained the following equation relating *r_50_* to *Q* and *K_M_* when corresponding kinetic rate parameters for the modifying and demodifying enzymes are identical (Supp Info Sec. 1.6):

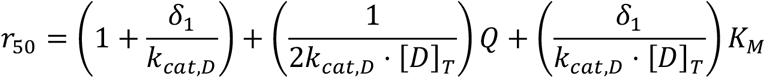

Considering the endpoints of the plots in Fig. 2c and the equation above, it is clear that as the rate of substrate production is made arbitrarily large, the *r*_50_ grows without bound. Thus the system described by the intermediate model becomes less and less sensitive to incoming signals. However, when we make *K_M_* as small as possible with all other parameters fixed (i.e. *K_M_* → 0), we see that 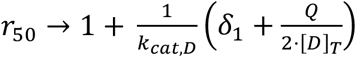, which is a constant. Note that this constant value is nevertheless larger than *r*_50_ = 1 for the GK model (Supp Info Sec. 1.6).

In a similar fashion we can analyze the effective Hill coefficient (*n_eff_*) for the intermediate model. Note that the *S\** vs. *r* curves do not precisely follow the form of a Hill function; as such, we use the standard definition 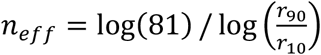 (6, 29). We can establish a lower bound on *n_eff_* for the Intermediate model (which we will refer to as *n_eff_*(I)), indicating that the intermediate model always exhibits positive cooperativity (i.e. *n_eff_*(I) > 1, Supp Info Sec. 1.7). As with *r*_50_, we also find that varying saturation by changing *Q* or *K_M_* results in opposing effects on *n_eff_* (Fig. 2d, Supp Info Figure S2). While *n_eff_*(GK) grows without bound in either case, *n_eff_*(I) is smaller by several orders of magnitude. For instance, when *Q* is increased, *n_eff_*(I) evaluates to exactly *2* regardless of the values of the other parameters.

The above results clearly demonstrate that changing saturation by varying *Q* and *K_M_* have very different consequences for the steady-state response of the PTM cycle in the Intermediate model. Since the value of *K_M_* depends on the underlying rate constants for the enzyme-substrate interaction, it is unlikely to vary on short time scales. As such, it is more likely that saturation will change by changing the production rate *Q in vivo*. Certainly, experimental manipulation of saturation generally occurs through changes in protein expression (e.g. by “overexpressing” the protein, which would correspond to increasing *Q* in this model). The steady-state responses of PTM cycles *in vivo* may thus be quite different from the standard predictions that have been made in the absence of any consideration of protein turnover (7–12).

### Driving protein degradation: “Full” model

While the results described above hold for any PTM cycle subject to turnover, we are ultimately interested in PTMs like ubiquitylation that drive protein degradation. This corresponds in our case to *δ*_2_ > *δ*_1_, which we term the “Full” model. For the purposes of display, we kept *δ*_1_ close to the average degradation rate of human proteins and set *δ*_2_ close to the fastest degradation rate observed in human cells (i.e. *δ*_1_ = 2 × 10^−5^ *s*^−1^ and *δ*_2_ = 2 × 10^−4^*s*^−1^) (27).

We first considered how changes in E3 ligase activity relative to DUB activity would influence the modification state of the substrate. We found that transitions in [*S*\*] are even less sensitive to incoming signals in the full model, compared to the intermediate model (Figs. 2a & 2b). Indeed, the *r_50_* for the full model is always greater than that for the intermediate model as *Q* is increased (Fig. 2c), and we have shown analytically that this is true for any reasonable set of kinetic parameters (Supp Info Sec. 1.6). Interestingly, although *n_eff_* (I) is always less than *n_eff_*(GK) as discussed above, we see that *n_eff_* (Full) is less than *n_eff_*(I) only for very small *Q* (Fig. 2d). For instance, when *Q* is increased without bound *n_eff_*(Full) evaluates to exactly 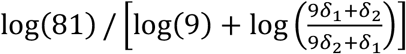, or approximately 7 in our case, which is larger than 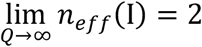.

Since E3 ligase activity drives higher levels of protein degradation in the full model, changes in the *r* parameter will change not only [*S\**] but also the total concentration of substrate ([*S*]*_T_*). Perhaps not surprisingly, we found that [*S*]*_T_* also exhibits an ultrasensitive transition in *r*. As with the transitions in [*S\**] discussed above, there is a rightward shift in r_50_ for the [*S*]*_T_* vs. *r* curve as *Q* is increased in the full model (Supp Info Sec. 1.8). This phenomenon can generate interesting behaviors, as shown in Fig. 3a. Suppose that we systematically increase the expression level of the protein while keeping the concentrations of the modifying/demodifying enzymes constant, which corresponds to a constant *r* in this model. As *Q* increases, the *r*_50_ of the curve increases from a point less than the value of *r* to a point greater than *r*. This leads to nonlinear changes in total substrate as Q increases (see below). The [*S*]*_T_* vs. *r* curve has a number of other similarities to the [*S\**] vs. *r* curve; for instance, we see that the effective Hill coefficient for total substrate in the full model does not change significantly when *Q* is increased, and never exceeds a value of 2 (Fig. 3b). In any case, this work clearly demonstrates that PTMs leading to increased rates of protein degradation can produce ultrasensitive transitions in total protein concentration (Figs. 2 and 3).

**Fig. 3.**
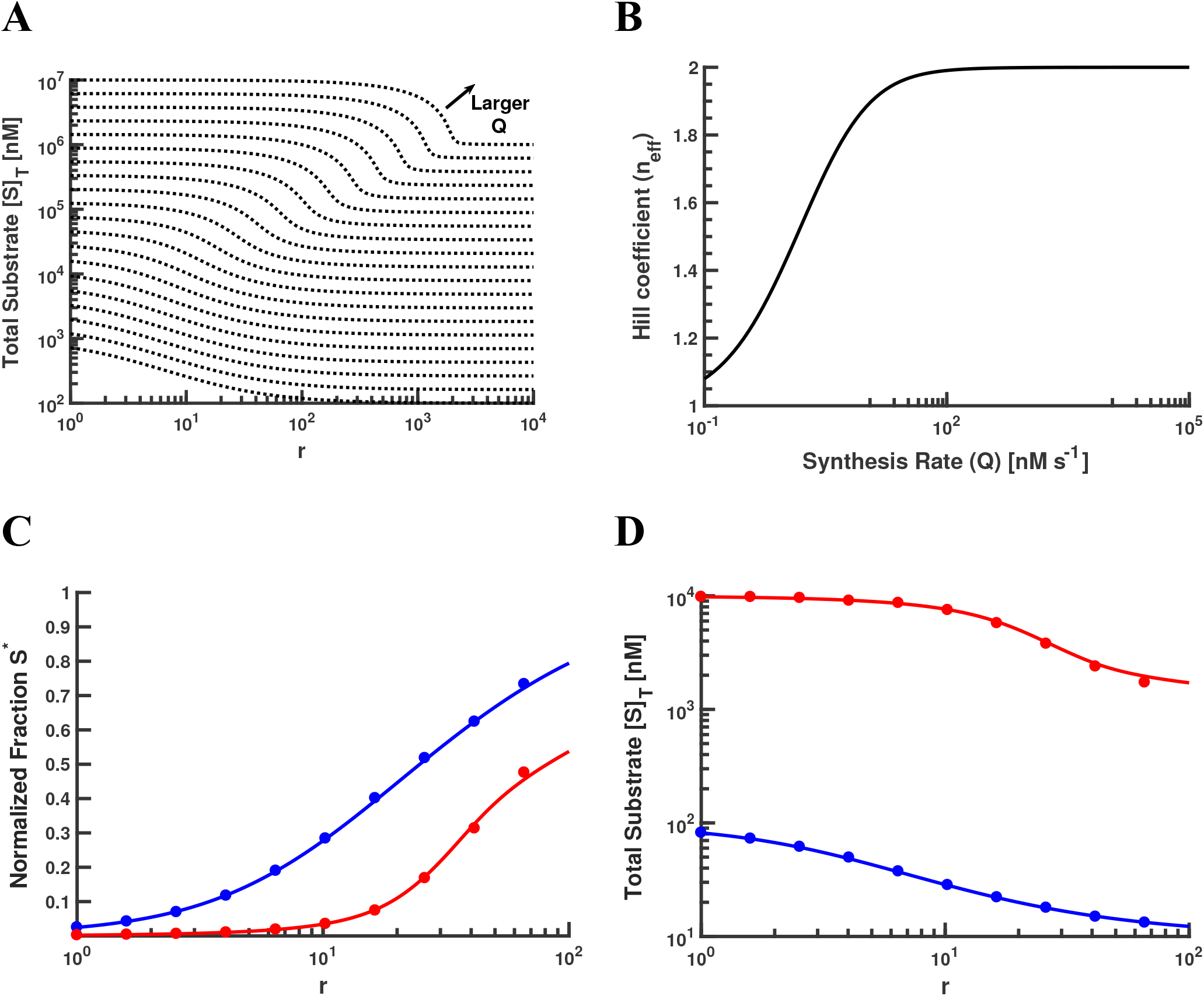
Effects of varying Q on the r_50_ and n_eff_ in the Full model and its representative analog for multiple modification states. **(A)** The shift in r_50_ for larger *Q* is clearly demarcated by the line in red. Each dashed curve indicates a different value for Q. The maximum and minimum values of [S]_T_ for each curve is Q/δ_1_ and Q/δ_2_, respectively. **(B)** The effective Hill coefficient n_eff_ is relatively unaffected by increase in protein synthesis rate, for the Full model. **(C)** In the Processive E3, Distributive/Sequential DUB model, much more E3 ligase activity is necessary for a maximal response in the saturated regime. The curves denote trajectories obtained by deterministic simulations, and the circles denote stochastic averages for the trajectories. There is very good agreement between both frameworks. **(D)** Compared to Panel (A), the rightward shift is more pronounced in the presence of polyubiquitin chains.

### Adding multiple modification states to the full model

While the full model is suggestive, it abstracts a number of details of the biological systems that control protein homeostasis. For instance, E3 ligases, rather than adding just a single ubiquitin to their substrates, instead tend to attach polyubiquitin chains of varying lengths (4). To capture the effects of this in our models, we surveyed available literature and found that multiple enzymatic mechanisms have been proposed for both E3 ligases and DUB enzymes (13, 30-38). E3 ligases may be “processive,” in the sense that the ligase adds an ubiquitin unit to the polyubiquitin chain at each catalytic step and stays attached to the substrate while multiple ubiquitins are added sequentially. Alternatively, they may be “distributive,” meaning that the ligase disassociates from the substrate at the end of each catalytic reaction. In the one case that has been extensively studied experimentally, a form of E3 called a RING ligase works with the E2 Cdc34 to build polyubiquitin chains on substrates in a processive manner (37). Of course, this does not mean that other E3 ligases might not display distributive kinetics. Regarding the DUB enzyme counterpart, 3 such enzymes have been found in 26S proteasomes: Rpn11, Usp14, and Uch37 (32–34, 38). Rpn11 functions by truncating at the base of the chain (in a distributive manner), whereas Usp14 and Uch37 serve primarily to trim the ubiquitin chains sequentially (in a processive manner). Interestingly, more than one DUB might act on a given chain (32).

Although there are experimentally characterized examples for several of these possible mechanisms, little is actually known about how widespread each mechanism may be in nature. We thus employed an exhaustive approach, examining all combinations of the enzyme mechanisms and creating models of those scenarios. Our parameter values for the distributive cases correspond to the values in the previous section (i.e. Single Substrate, Single Modification State). However, we used parameter values directly from literature for the processive cases (37).

Given available experimental data (37), we focus on a reasonable and representative model (i.e. Processive E3 and Distributive/Sequential DUB) from this set of models. For purposes of display, we have depicted this scheme in Fig. 1c. It is known that a polyubiquitin chain typically requires at least four ubiquitin units to be effectively degraded by the proteasome (13, 30, 31). We thus assumed that each of the first three modification states (0-3 ubiquitins) is degraded at a uniform rate, *δ*_1_, which is smaller than the corresponding (higher) rate *δ*_2_ for each of the remaining states (4 or more ubiquitins).

In theory, the ubiquitin chain could reach an infinite length, though of course in practice the action of DUBs and degradation will limit the largest chain typically observed in the system. Denoting this maximum length of the ubiquitin chain by *ℓ*, we enumerated the chemical reaction networks for all of the possible mechanistic scenarios described above (Supp Info Sec. 2.1). Due to the inherent complexity of the model, we could not obtain closed-form analytical solutions, and thus focused on numerical simulations.

Recall that in the full model, which has a single modification state, we found a significant reduction in both sensitivity and ultrasensitivity of the transition in *S\** when compared to the intermediate model. To compare our more complex model with the full model, we defined *S\** for the case with multiple modification states as follows: 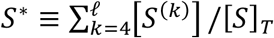, where *k* indexes the substrate modification state. To choose a reasonable value for *ℓ*, we systematically increased this parameter and found a threshold value such that changes in *r*_50_ and *n_eff_* were negligible beyond that threshold. Using this approach, we chose a value of 500 for *ℓ* heuristically by visual inspection. To investigate the effects of allowing for arbitrarily large chain length, we also performed simulations using an agent-based stochastic modeling framework (39–42). In Figs. 3c & 3d, we see that the inclusion of ubiquitin chains magnifies the aforementioned effects in both the *S\** vs. *r* and [*S*]*_T_* vs. *r* curves. Specifically, much more E3 ligase activity is necessary to achieve a maximal response in saturated regimes. Furthermore, there is excellent agreement between the deterministic framework and its stochastic counterpart, suggesting that truncating the system at *ℓ* = 500 yields a reasonable approximation. All of the models that we examined, arising from the various mechanisms proposed for the E3 ligase and DUB enzymes, generated similar qualitative behavior (Supp Info Sec. 2.2), which indicates that these findings are largely invariant with respect to the catalytic mechanisms utilized by the E3 ligase and DUB enzymes (4).

### Adding multiple substrates to the full model

As mentioned above (Fig. 3a), there is an increase in *r*_50_ for the [*S*]*_T_* vs. *r* curve as *Q* is increased in the full model. As a consequence, increasing *Q* while keeping *r* fixed results in the curve seen in Fig. 4a. For low values of *Q*, the transition in *r*_50_ occurs before this fixed *r*-value, so [*S*]*_T_* ≈ *Q/δ*_2_; for large Q, the transition in *r*_50_ occurs after this fixed *r*-value, so [*S*]*_T_* ≈ *Q/δ*_1_. For intermediate *Q*, however, there is a distinct transition between these two regimes. The result in Fig. 4a implies that if two substrates share an E3/DUB enzyme pair, the *r*_50_’s of the transitions in total substrate concentrations for these two proteins will be coupled. Thus, overexpressing one protein might have an influence on the expression profile of its counterpart.

**Fig. 4.**
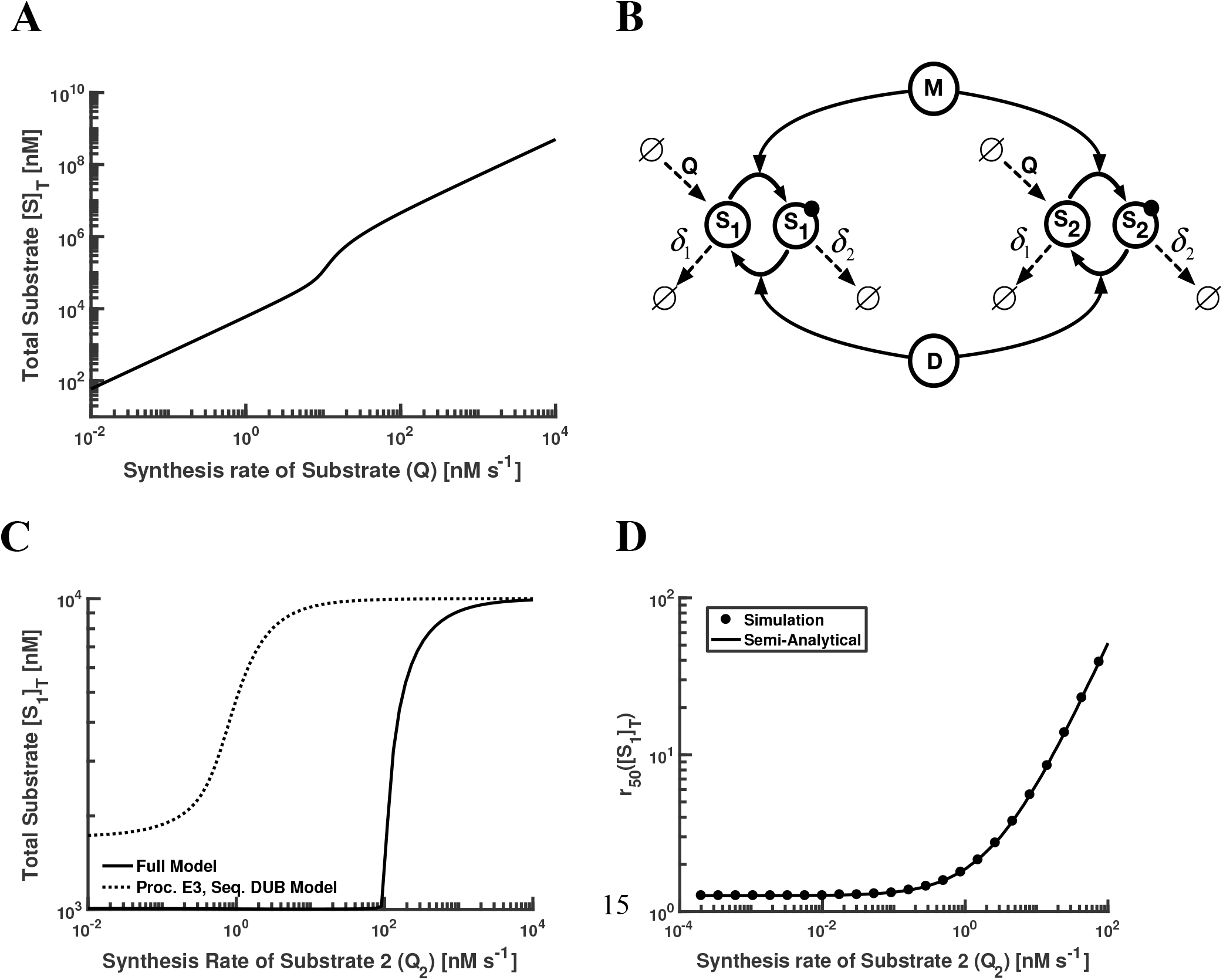
Effects of protein overexpression on total protein in single-substrate and multiple-substrate models. **(A)** There is a qualitative transition in the [S]_T_ vs. *Q* curve when *Q* is comparable in magnitude with the measure of saturation (in this case, [M]). The slope of the curve approaches 1 on either side of the transition, which is consistent with the maximum and minimum values of [S]_T_. **(B)** Schematic diagram for multiple substrates with one modification state each. Shown here is the model corresponding to two substrates, for simplicity. Here “M” denotes modifying enzyme and “D” denotes demodifying enzyme. Modified substrate is indicated by the dark circle. **(C)** Sensitivity to signal for first substrate vs. synthesis rate of the second substrate, for the multiple-substrate analog of the Full model. The semi-analytical curve was obtained by substituting certain values, obtained empirically from simulation, into the analytical expression for r_50_([S_1_]_T_). Axes in log scale. **(D)** Plot of total concentration of first substrate vs. synthesis rate of the second substrate, for the multiple-substrate analogs of the Full model and the representative model. Axes in log scale.

To test this, we introduced more than one substrate in the context of the full model. For the sake of display, we have taken the total number of substrates *N* in our model to be 2. As shown in Fig. 4b, each E3 ligase and DUB now acts on two substrates with one modification state per substrate. The set of ODEs describing the model is given in Supp Info Sec. 3. In contrast to the case of just one substrate, here we are interested in capturing the response of 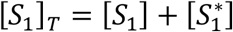 to changes in the synthesis rate of the second substrate, denoted by *Q*_2_. In Fig. 4c, we see that increasing *Q*_2_ yields an increase in [*S*_1_]*_T_* for the Full model, as expected. The general behavior is similar in the Processive E3 and Distributive/Sequential DUB model, with the transition in [*S*_1_]*_T_* occurring at lower values of *Q*_2_.

Interestingly, *r*_50_([*S*_1_]*_T_*) also depends on *Q*_2_ (Fig. 4d). Specifically, when *Q*_2_ is large enough in the Full model, *r*_50_ increases linearly with respect to *Q*_2_. In a similar manner to Fig. 4c, a lower value of *Q_2_* is sufficient to obtain a linear increase in *r*_50_ in the presence of multiple modification states. Note the excellent agreement in the Full model between the *r*_50_ values extracted empirically from simulation output and the analytical expression for *r*_50_([*S*_1_]*_T_*) (Fig. 4d & Supp Info Sec 3.3). In fact, when we make *Q*_2_ arbitrarily large, we obtain

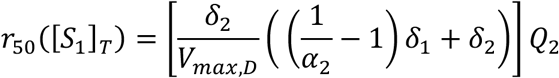

where *α*_2_ is the molar fraction of modified *S*_2_ at steady-state when [*S*_1_]*_T_* is half-maximal. In other words, as the concentration of the second substrate is increased, it takes more and more activity of the E3 ligase to drive the transition in *S*_1_ concentration. Interestingly, all of these results can be readily generalized for any number of substrates, independent of substrate identity (Supp Info Sec. 3). Thus, crosstalk in PTMs can lead to coupling of not only modification states (18, 19), but also of overall protein levels.

## Discussion

It has been over 35 years since Goldbeter and Koshland discovered the phenomenon of 0^th^-order ultrasensitivity. Since then, there has been extensive characterization of PTM cycles with 0^th^-order ultrasensitivity, both experimentally (43–45) and computationally (18, 20, 25, 46, 47). Until now, however, the properties of PTM cycles that drive protein degradation have not been studied in a systematic way. Using a mathematical modeling framework, we found that adding synthesis and degradation to a PTM cycle suppresses both sensitivity to signal and ultrasensitivity of the response, even when the PTM in question does not serve as a signal for protein degradation. Thus switch-like behaviors *in vivo* may or may not be the consequence of 0^th^-order ultrasensitivity, depending on the stability of the protein substrate. Although there are exceptions (15–17), most models of signaling networks ignore protein turnover (48, 49). Our findings indicate that incorporating turnover, especially turnover based on actual protein stabilities, is key to capturing the global PTM dynamics of signaling systems.

Interestingly, we found the general trend of decreasing sensitivity and ultrasensitivity holds for PTMs that drive protein degradation, even when accounting for many of the complicated mechanisms that describe polyubiquitylation by E3 ligases and deubiquitylation by DUB enzymes (Fig. 3 & Supp Info Sec. 2). By adding E3 ligase crosstalk, we demonstrated that overexpressing one protein can elevate the concentration of another, and can also reduce the sensitivity of other proteins to incoming signals that would drive their degradation (Figs. 4c, 4d). In other words, if one protein is overexpressed, it becomes more difficult to degrade any of its counterparts sharing the same E3/DUB enzyme pair.

Although there is some data available about the specificity of E3 ligases (34, 50, 51), this information is very far from complete. Consider the highly common experimental scenario where a primary aim is to characterize the function of a protein by manipulating its expression level. Our findings indicate that the interpretation of overexpression data in eukaryotic cells may be very difficult because some of the observed phenotypic or molecular effects could be directly due to the higher concentration of the protein that was expressed, but other effects could be due to E3 ligase coupling (Fig. 4c). Additional complications could also appear due to the change in sensitivity to the shared E3 ligases for other substrates in the system (Fig. 4d). For instance, if a protein is being actively regulated by its E3 ligase and a degradation signal appears, then a high concentration of other proteins in the system would potentially inhibit the signal. This could have unforeseen large-scale effects on the overall system.

A global picture of E3-ligase/DUB enzyme specificity will thus likely be essential to comprehending the regulation of protein levels within cells. This will allow us to begin determining how to isolate direct effects of changes in protein expression levels from indirect effects. Equally necessary are mathematical or computational models of signaling dynamics, gene regulatory networks, and other cellular processes that describe the interplay between PTMs that do not lead to degradation and those that drive degradation. Incorporating the coupled dynamics of protein levels into our understanding of cell signaling and cellular physiology thus represents a grand challenge for both experimental and computational systems biology.

## Materials and Methods

**Experimental methods:** Our model behaviors can be described deterministically by systems of ordinary differential equations (ODEs). Numerical integration of the systems was performed by the stiff solver ode15s in MATLAB. All analyses were performed at steady-state. In parallel, agent-based stochastic simulations of the systems (39–41) were conducted using custom-built software implemented in C++. Parameter values were chosen to ensure equivalence between the deterministic and stochastic systems. See the supporting information for full details regarding all the models considered here. All simulation software codes are available upon author request.

## Acknowledgements

We thank William Mather and Christian Ray for helpful discussions regarding this work.

