## Supporting Information for "Crosstalk and Ultrasensitivity in Protein Degradation Pathways"

Abhishek Mallela[1], Maulik K. Nariya[2] and Eric J. Deeds[3,4]

[1]Department of Mathematics, University of California Davis, Davis, CA, USA
[2]Laboratory of Systems Pharmacology, Harvard Medical School, Boston, MA, USA
[3]Department of Integrative Biology and Physiology, University of California Los Angeles, Los Angeles, CA, USA
[4]Institute for Quantitative and Computational Biosciences, University of California Los Angeles, Los Angeles, CA, USA

**4 Multiple Substrates, Multiple Modification States** **23**

**References** **23**

### 1 Single Substrate, Single Modification State

We begin with a modification of the Goldbeter-Koshland loop that incorporates protein turnover. It is described by the following schematic of enzymatic reactions:

$$M + S \underset{k_{-,M}}{\overset{k_{+,M}}{\rightleftharpoons}} MS \xrightarrow{k_{cat,M}} M + S^*$$

$$D + S^* \underset{k_{-,D}}{\overset{k_{+,D}}{\rightleftharpoons}} DS^* \xrightarrow{k_{cat,D}} D + S$$

$$\emptyset \xrightarrow{Q} S$$

$$S \xrightarrow{\delta_1} \emptyset$$

$$S^* \xrightarrow{\delta_2} \emptyset$$

$$MS \xrightarrow{\delta_1} M$$

$$DS^* \xrightarrow{\delta_2} D$$

Here $M$ and $D$ represent any modifying and demodifying enzyme, respectively. Any substrate in the $S$ state is degraded at a first-order rate $\delta_1$, and any in the $S^*$ state at a rate $\delta_2$, with $\delta_2 > \delta_1$. Substrate is also synthesized at a constant rate $Q$, and all the synthesized substrates are in the $S$ (unmodified) state. Note that degradation does not consume $M$ or $D$.

It is straightforward to use the Law of Mass Action to formulate the corresponding set of ordinary differential equations (ODEs), as described below.

#### 1.1 Equations

##### *1.1.1 ODEs*

$$\frac{d[S]}{dt} = Q - k_{+,M}[M][S] + k_{-,M}[MS] + k_{cat,D}[DS^*] - \delta_1[S] \tag{1}$$

$$\frac{d[S^*]}{dt} = -k_{+,D}[D][S^*] + k_{-,D}[DS^*] + k_{cat,M}[MS] - \delta_2[S^*] \tag{2}$$

$$\frac{d[MS]}{dt} = k_{+,M}[M][S] - (k_{-,M} + k_{cat,M} + \delta_1)[MS] \tag{3}$$

$$\frac{d[DS^*]}{dt} = k_{+,D}[D][S^*] - (k_{-,D} + k_{cat,D} + \delta_2)[DS^*] \tag{4}$$

$$\frac{d[M]}{dt} = (k_{-,M} + k_{cat,M} + \delta_1)[MS] - k_{+,M}[M][S] \tag{5}$$

$$\frac{d[D]}{dt} = (k_{-,D} + k_{cat,D} + \delta_2)[DS^*] - k_{+,D}[D][S^*] \tag{6}$$

##### 1.1.2 Mass conservation

$$[M]_T = [M] + [MS]$$
$$[D]_T = [D] + [DS^*]$$
$$[S]_T = [S] + [S^*] + [MS] + [DS^*]$$

#### 1.2 Model definitions

Using the formalism above, we can describe the Goldbeter-Koshland model as having no synthesis or degradation (i.e. with $Q = \delta_2 = \delta_1 = 0$). The Intermediate model has a positive synthesis rate and a uniform rate of degradation (i.e. with $Q > 0$ and $\delta_2 = \delta_1 > 0$.) Finally, the Full model also has a positive synthesis rate, but has different rates of degradation (i.e. $Q > 0$ and $\delta_2 > \delta_1 > 0$.)

#### 1.3 Preliminaries

##### 1.3.1 Steady-state

Let $K_{M,1} \equiv \frac{k_{-,M}+k_{cat,M}+\delta_1}{k_{+,M}} = K_{M,M} + \frac{\delta_1}{k_{+,M}}$ and $K_{M,2} \equiv \frac{k_{-,D}+k_{cat,D}+\delta_2}{k_{+,D}} = K_{M,D} + \frac{\delta_2}{k_{+,D}}$. We study the system at steady-state (i.e. setting the L.H.S. of eq. (1) - eq. (6) equal to zero). From the steady-state versions of eq. (3) and eq. (4), $[MS] = \frac{[M]\cdot[S]}{K_{M,M}}$ and $[DS^*] = \frac{[D]\cdot[S^*]}{K_{M,D}}$. Then

$$[M]_T = [M] + [MS] = [M]\left(1 + \frac{[S]}{K_{M,M}}\right) \implies [MS] = \frac{\left(\frac{[M]_T}{1+\frac{[S]}{K_{M,M}}}\right)[S]}{K_{M,M}} = \frac{[M]_T[S]}{K_{M,M} + [S]}$$

$$[D]_T = [D] + [DS^*] = [D]\left(1 + \frac{[S^*]}{K_{M,D}}\right) \implies [DS^*] = \frac{\left(\frac{[D]_T}{1+\frac{[S^*]}{K_{M,D}}}\right)[S^*]}{K_{M,D}} = \frac{[D]_T[S^*]}{K_{M,D} + [S^*]}$$

Adding eq. (1) and eq. (3) at steady-state yields

$$Q - \delta_1[S] + k_{cat,D}[DS^*] = (k_{cat,M} + \delta_1)[MS] \quad (7)$$

Adding eq. (2), eq. (4) at steady-state gives

$$k_{cat,M}[MS] - \delta_2[S^*] = (k_{cat,D} + \delta_2)[DS^*] \quad (8)$$

##### 1.3.2 Equivalence

In order to proceed with our analysis, we first establish that eq. (7) is equivalent to eq. (8). Note that adding eq. (1) - eq. (4) at steady-state yields

$$Q = \delta_1([S] + [MS]) + \delta_2([S^*] + [DS^*]) \tag{9}$$

Substituting eq. (9) in eq. (7) gives

$$\delta_1[MS] + \delta_2[S^*] + \delta_2[DS^*] + k_{cat,D}[DS^*] = (k_{cat,M} + \delta_1)[MS]$$

which is equivalent to eq. (8) upon rearrangement.

##### 1.3.3 Total substrate

By the mass conservation equations, since $[M], [D] \geq 0$, we have $[MS] \leq [M]_T$ and $[DS^*] \leq [D]_T$. In the derivations that follow, we make the standard (Michaelis-Menten) assumption that total substrate is much larger than the concentration of either enzyme, so $[S]_T \approx [S] + [S^*]$. Note that eq. (9) implies $[S]_T = \frac{Q}{\delta_1}$ in the Intermediate model, since $\delta_1 = \delta_2$. For the full model, eq. (9) gives:

$$\begin{aligned} Q = \delta_1([S] + [MS]) + (\delta_1 + \delta_2 - \delta_1)([S^*] + [DS^*]) &\implies Q = \delta_1[S]_T + (\delta_2 - \delta_1)([S^*] + [DS^*]) \\ &\implies [S]_T \approx \frac{Q}{\delta_1} + \left(1 - \frac{\delta_2}{\delta_1}\right)[S^*] \end{aligned}$$

For purposes of display, the case of equal $K_M$'s (i.e. $K_{M,M} = K_{M,D}$) is analyzed below. Analyses of scenarios with substantially different $K_M$s are left to future work. It follows that $K_{M,1} \approx K_{M,2}$ because $k_{+,M}, k_{+,D} \gg \delta_1, \delta_2$ for the majority of enzymes.

##### 1.3.4 Parameter values

The following figure validates the biological relevance of our choices of parameter values. The curve in each plot is a kernel density estimate of the experimental values obtained from the BRENDA enzyme database.

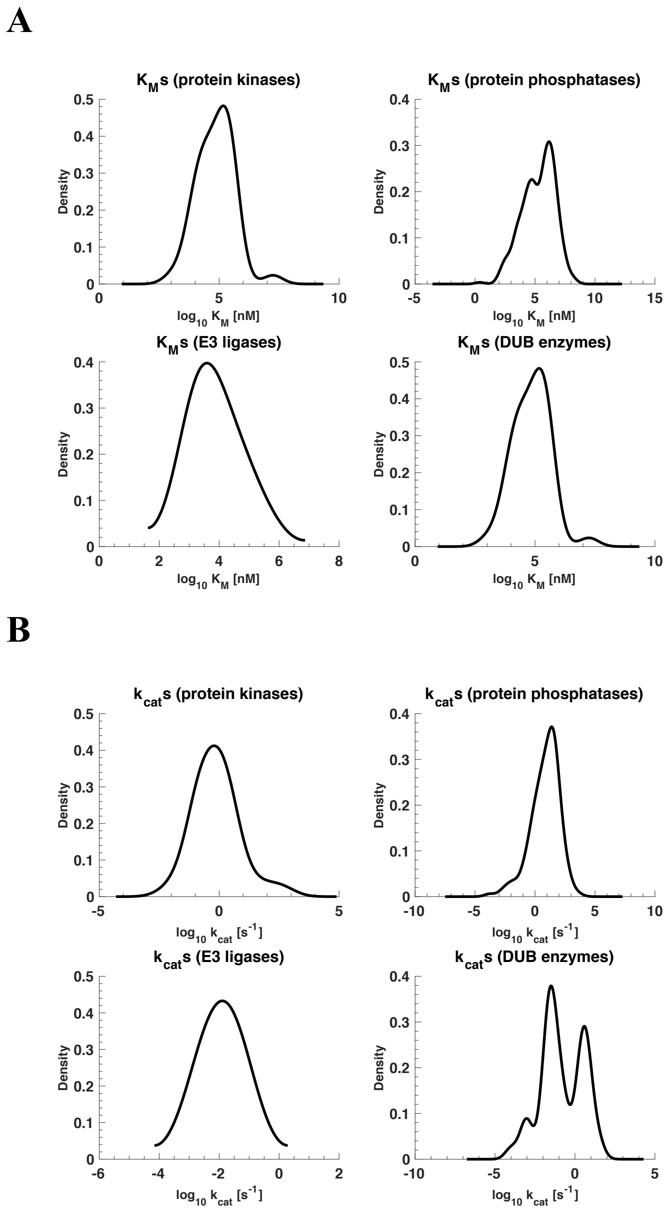

Figure S1: **Distributions of relevant steady-state kinetic parameters.** (A) Logarithmic-scale density plots of the $K_M$s for protein kinases, protein phosphatases, E3 ligases, and DUB enzymes in the human genome. Experimental values were obtained from the BRENDA enzyme database (1) by using EC numbers for the tyrosine kinases/phosphatases and serine/threonine kinases/phosphatases. (B) Identical procedure as in panel (A); implemented for $k_{cat}$s.

Table S1: **Parameter values used for Figs. 2a, 2b in the main text.** For Fig. 2a: chosen on the basis of $\delta_1 = \log(2)/(10 \text{ hours})$, which is the average reported protein half-life in human cells; for Fig. 2b: $K_M = 10^1 \times [S]_T$ (unsaturated) and $K_M = 10^{-1} \times [S]_T$ (saturated)

| | **Fig. 2a** | **Fig. 2b** | **Units** |
|---|---|---|---|
| $\boldsymbol{Q(unsat.)}$ | $2 \times 10^{-2}$ | $2 \times 10^{-2}$ | $[nM] \cdot [s]^{-1}$ |
| $\boldsymbol{Q(sat.)}$ | 2 | $2 \times 10^{-2}$ | $[nM] \cdot [s]^{-1}$ |
| $\boldsymbol{k_+(unsat.)}$ | $1 \times 10^{-4}$ | $1 \times 10^{-4}$ | $[nM \cdot s]^{-1}$ |
| $\boldsymbol{k_+(sat.)}$ | $1 \times 10^{-4}$ | $1 \times 10^{-2}$ | $[nM \cdot s]^{-1}$ |
| $\boldsymbol{k_-}$ | $1 \times 10^{-3}$ | $1 \times 10^{-3}$ | $[s]^{-1}$ |
| $\boldsymbol{k_{cat}}$ | 0.999 | 0.999 | $[s]^{-1}$ |
| $\boldsymbol{\delta_1}$ | $2 \times 10^{-5}$ | $2 \times 10^{-5}$ | $[s]^{-1}$ |
| $\boldsymbol{\delta_2}$ | $2 \times 10^{-4}$ | $2 \times 10^{-4}$ | $[s]^{-1}$ |

#### 1.4 Analytical expressions for $r_{50}$

Let $r$ denote the ratio of maximum velocities of the two enzymes (i.e. $r \equiv \frac{V_{max,M}}{V_{max,D}} = \frac{k_{cat,M}[M]_T}{k_{cat,D}[D]_T}$). This ratio represents the signal source for the system as defined previously (2). Let $\alpha \equiv [S^*]/[S]_T$, or the molar fraction of modified substrate at steady-state. The $r_{50}$ of the response is defined as the amount of $r$ necessary to yield a 50% response in modified substrate, or $\alpha = 0.5$.

We can rewrite eq. (8) by replacing $[MS]$ and $[DS^*]$ with their Michaelis-Menten forms. We can then divide through by $k_{cat,D}[D]_T$ and the resulting equation can be solved for $r$:

$$\begin{aligned}
\frac{k_{cat,M}[M]_T[S]}{K_M + [S]} - \delta_2[S^*] &= [S^*]\left(\frac{k_{cat,D}[D]_T}{K_M + [S^*]} + \frac{\delta_2[D]_T}{K_M + [S^*]}\right) \\
\frac{k_{cat,M}[M]_T[S]}{k_{cat,D}[D]_T(K_M + [S])} - \frac{\delta_2[S^*]}{k_{cat,D}[D]_T} &= [S^*]\left(\frac{1}{K_M + [S^*]} + \frac{\delta_2[D]_T}{(k_{cat,D}[D]_T)(K_M + [S^*])}\right) \\
\frac{r[S]}{K_M + [S]} &= \frac{[S^*]}{k_{cat,D}}\left(\frac{\delta_2}{[D]_T} + \frac{\delta_2 + k_{cat,D}}{K_M + [S^*]}\right) \\
r &= \frac{[S^*]}{k_{cat,D}}\left(\frac{\delta_2}{[D]_T} + \frac{\delta_2 + k_{cat,D}}{K_M + [S^*]}\right)\left(1 + \frac{K_M}{[S]}\right) \qquad (10)
\end{aligned}$$

##### *1.4.1 Intermediate Model*

For the intermediate model, since $Q, \delta_1 > 0$, note that eq. (10) becomes:

$$\begin{aligned}
r &= \frac{\alpha}{k_{cat,D}}\left(\frac{\delta_1[S]_T}{[D]_T} + \frac{(\delta_1 + k_{cat,D})[S]_T}{K_M + [S^*]}\right)\left(1 + \frac{K_M}{(1-\alpha)[S]_T}\right) \\
&= \frac{\alpha}{k_{cat,D}}\left(\frac{Q}{[D]_T} + \frac{Q(\delta_1 + k_{cat,D})}{\delta_1 K_M + \delta_1[S^*]}\right)\left(1 + \frac{\delta_1 K_M}{(1-\alpha)Q}\right) \\
&= \frac{Q\alpha}{k_{cat,D}}\left(\frac{1}{[D]_T} + \frac{(\delta_1 + k_{cat,D})}{\delta_1 K_M + \alpha Q}\right)\left(1 + \frac{\delta_1 K_M}{(1-\alpha)Q}\right) \qquad (11)
\end{aligned}$$

We can substitute $\alpha = 0.5$ in the expression above to obtain $r_{50}$. After simplification (3), we obtain

$$r_{50} = 1 + \frac{Q}{2k_{cat,D}[D]_T} + \frac{\delta_1(K_M + [D]_T)}{k_{cat,D}[D]_T} \tag{12}$$

##### 1.4.2 Full Model

For the full model, we directly solve for $[S], [S^*]$ in terms of $\alpha$. Observe that $[S^*] = \alpha[S]_T = \frac{\alpha Q}{\delta_1} + \alpha[S^*](1 - \frac{\delta_2}{\delta_1})$. Solving for $[S^*]$ we get

$$[S^*] = \frac{\alpha Q}{\delta_1 + \alpha(\delta_2 - \delta_1)}.$$

Since $[S] = [S]_T - [S^*] = \frac{(Q - \delta_2[S^*])}{\delta_1}$, we have

$$[S] = \frac{(1-\alpha)Q}{\delta_1 + \alpha(\delta_2 - \delta_1)}.$$

Plugging these expressions into eq. (10), we arrive at a closed-form expression for $r$ as a function of $\alpha$:

$$r = \frac{\alpha Q}{k_{cat,D}\left[\delta_1 + \alpha(\delta_2 - \delta_1)\right]} \left( \frac{\delta_2}{[D]_T} + \frac{\delta_2 + k_{cat,D}}{K_M + \frac{\alpha Q}{\delta_1 + \alpha(\delta_2 - \delta_1)}} \right) \left( 1 + \frac{K_M}{\frac{(1-\alpha)Q}{\delta_1 + \alpha(\delta_2 - \delta_1)}} \right) \tag{13}$$

Substituting $\alpha = 0.5$ in eq. (13) and simplifying the resulting expression (3) yields:

$$r_{50} = 1 + \frac{Q\delta_2}{k_{cat,D}[D]_T(\delta_1 + \delta_2)} + \frac{\delta_2(K_M + [D]_T)}{k_{cat,D}[D]_T} \tag{14}$$

#### 1.5 Analytical expressions for $n_{eff}$

The effective Hill coefficient $n_{eff}$ is defined in (4) as $\log(81)/\log(\frac{EC_{90}}{EC_{10}})$. From (5), we have the expression for $n_{eff}$ in the Goldbeter-Koshland model:

$$n_{eff}(\text{GK}) = \frac{\log(81)}{\log(81) + 2\log\left( \frac{\frac{K_M}{[S]_T} + 0.1}{\frac{K_M}{[S]_T} + 0.9} \right)}$$

We can substitute $\alpha = 0.1, 0.9$ in both eq. (11) and eq. (13) to yield both $EC_{10}$ and $EC_{90}$ for the intermediate and full models, respectively. Simplifying, we obtain:

$$n_{eff}(\text{Intermediate}) = \frac{\log(81)}{\log(81) + 2\log\left(\frac{\frac{K_M}{[S]_T} + 0.1}{\frac{K_M}{[S]_T} + 0.9}\right) + \log\left(\frac{\frac{(k_{cat,D}+\delta_1)[D]_T}{Q} + \frac{K_M}{[S]_T} + 0.9}{\frac{(k_{cat,D}+\delta_1)[D]_T}{Q} + \frac{K_M}{[S]_T} + 0.1}\right)}$$

and

$$n_{eff}(\text{Full}) = \frac{\log(81)}{\log(81) + \log\left(\frac{x(Q+K_Mx)(Q+K_My)(\delta_2[9Q + y(K_M + [D]_T)] + [D]_Tk_{cat,D}y)}{y(9Q+K_Mx)(9Q+K_My)(\delta_2[Q + x(K_M + [D]_T)] + [D]_Tk_{cat,D}x)}\right)}$$

where $x = 9\delta_1 + \delta_2$ and $y = 9\delta_2 + \delta_1$.

##### 1.6 Analysis of $r_{50}$ in saturated regimes

We show that $r_{50}(\text{GK}) < r_{50}(\text{Intermediate}) < r_{50}(\text{Full})$ when total substrate is at saturating concentrations (i.e. when $[S]_T \gg K_M$).
First observe that $r_{50}(\text{GK}) = 1$ in a saturated regime. This can be seen by setting $\delta_2 = 0$ in eq. (10) and noting that $[S], [S^*] \gg K_M$:

$$r = \frac{[S^*]}{k_{cat,D}}\left(\frac{k_{cat,D}}{K_M + [S^*]}\right)\left(1 + \frac{K_M}{[S]}\right) = \frac{\frac{[S^*]}{K_M+[S^*]}}{\frac{[S]}{K_M+[S]}} = 1$$

Now

$$r_{50}(\text{GK}) < 1 + \frac{Q}{2k_{cat,D}[D]_T} + \frac{\delta_1(K_M + [D]_T)}{k_{cat,D}[D]_T} = \underbrace{1 + \frac{Q\delta_2}{k_{cat,D}[D]_T(\delta_2 + \delta_2)} + \frac{\delta_1(K_M + [D]_T)}{k_{cat,D}[D]_T}}_{r_{50}(\text{Intermediate})}$$

$$< \underbrace{1 + \frac{Q\delta_2}{k_{cat,D}[D]_T(\delta_1 + \delta_2)} + \frac{\delta_2(K_M + [D]_T)}{k_{cat,D}[D]_T}}_{r_{50}(\text{Full})}$$

Furthermore, we can show that different modes of enzyme saturation (i.e. increasing $Q$ vs. decreasing $K_M$) effect substrate responses in distinct ways. Specifically, varying $Q$ has a stronger effect on $r_{50}$ than varying $K_M$ - this is true in both the Intermediate and Full models. We use the following chain of (unitless) inequalities to order the quantities of interest:

$$\frac{Q\delta_2}{k_{cat,D}[D]_T(\delta_1 + \delta_2)} > \frac{Q\delta_2}{k_{cat,D}[D]_T} > \frac{K_M\delta_2}{k_{cat,D}[D]_T} > \frac{K_M\delta_1}{k_{cat,D}[D]_T}$$

which is equivalent to

$$\frac{\partial r_{50}(\text{Full})}{\partial \log Q} > \frac{\partial r_{50}(\text{Intermediate})}{\partial \log Q} > \frac{\partial r_{50}(\text{Full})}{\partial \log K_M} > \frac{\partial r_{50}(\text{Intermediate})}{\partial \log K_M}$$

The only condition on this chain is $\frac{Q}{\delta_1} > 2K_M$, which is trivially satisfied in the saturated regime (i.e. $[S]_T \gg K_M$).

##### 1.7 Analysis of $n_{eff}$

When total substrate is at saturating concentrations, since $n_{eff}$(Intermediate) has an additional positive term in the denominator compared to $n_{eff}$(GK), $n_{eff}$(Intermediate) $< n_{eff}$(GK). Although it would be desirable to compare $n_{eff}$(GK) with $n_{eff}$(Full), this is hard to do for two reasons. Firstly, $[S]_T$ is not a function of $r$ in the Goldbeter-Koshland model but $[S]_T$ is a function of $r$ in the Full model. Secondly, $[S]_T$ is the solution of a cubic equation in both models, making analysis difficult.

However, we can show that in the limit $K_M \to 0, n_{eff}$(Intermediate) is a strictly decreasing function in $Q$:

$$n_{eff}(\text{Intermediate}) = \frac{\log(81)}{\log(81) + \log\left(\frac{\frac{(k_{cat,D}+\delta_1)[D]_T}{Q} + 0.9}{\frac{(k_{cat,D}+\delta_1)[D]_T}{Q} + 0.1}\right)}$$

has a negative partial derivative with respect to $Q$. Entering the expression into Mathematica and simplifying yields:

$$\begin{aligned}\frac{\partial n_{eff}(\text{Intermediate})}{\partial Q} &= -\frac{0.8(k_{cat,D}+\delta_1)}{(k_{cat,D}+\delta_1+0.1Q)(k_{cat,D}+\delta_1+0.9Q)\left[\log(81) + \log\left(\frac{\frac{(k_{cat,D}+\delta_1)[D]_T}{Q} + 0.9}{\frac{(k_{cat,D}+\delta_1)[D]_T}{Q} + 0.1}\right)\right]^2} \\ &< 0\end{aligned}$$

We can also show that $n_{eff}$(Intermediate) $> 1$ for all possible values of the parameters in the model. This is equivalent to showing

$$\left(\frac{\frac{K_M}{[S]_T} + 0.1}{\frac{K_M}{[S]_T} + 0.9}\right)^2 \left(\frac{\frac{(k_{cat,D}+\delta_1)[D]_T}{Q} + \frac{K_M}{[S]_T} + 0.9}{\frac{(k_{cat,D}+\delta_1)[D]_T}{Q} + \frac{K_M}{[S]_T} + 0.1}\right) < 1$$

or

$$\left(\frac{10\delta_1 K_M + Q}{10\delta_1 K_M + 9Q}\right)^2 \left(\frac{10(k_{cat,D}+\delta_1)[P]_T + 10\delta_1 K_M + 9Q}{10(k_{cat,D}+\delta_1)[P]_T + 10\delta_1 K_M + Q}\right) < 1$$

or

$$(10\delta_1 K_M + Q)^2(10(k_{cat,D} + \delta_1)[P]_T + 10\delta_1 K_M + 9Q) <$$
$$(10\delta_1 K_M + 9Q)^2(10(k_{cat,D} + \delta_1)[P]_T + 10\delta_1 K_M + Q)$$

We can rewrite the above inequality as follows (3):

$$8Q[(10\delta_1 K_M + Q)^2 + 8Q(10\delta_1 K_M + Q) + 2(10\delta_1 K_M + Q)(10(k_{cat,D} + \delta_1)[P]_T) + 8Q(10(k_{cat,D} + \delta_1)[P]_T)] > 0$$

which is true because the expression on the left-hand side is a combination of products and sums of positive terms.

#### 1.8 Analytical expression for $r_{50}$ of $[S]_T$ in Full model

Defining $r$ as in the previous section (i.e. $r = \frac{k_{cat,M}[M]_T}{k_{cat,D}[D]_T}$), let $\beta \equiv [S]_T$, the amount of total substrate at steady-state. The $r_{50}$ of the response $\beta$ is defined as the amount of $r$ necessary to yield a 50% response in total substrate. This quantity is of interest only in the full model, since $\beta$ is not constant with respect to $r$.

The half-maximal response for total substrate occurs at $\frac{1}{2}(\min [S]_T + \max [S]_T)$. Since

$$[S]_T = \frac{Q}{\delta_1} + \left(1 - \frac{\delta_2}{\delta_1}\right)[S^*] \tag{15}$$

we can also solve for $[S]_T$ in terms of $[S]$:

$$[S]_T = \frac{Q}{\delta_1} + \left(1 - \frac{\delta_2}{\delta_1}\right)[S^*] \implies [S]_T = \frac{Q}{\delta_1} + \left(1 - \frac{\delta_2}{\delta_1}\right)([S]_T - [S])$$

Hence

$$[S]_T = \frac{Q}{\delta_2} + \left(1 - \frac{\delta_1}{\delta_2}\right)[S] \tag{16}$$

From eq. (15), we see that $[S]_T \leq \frac{Q}{\delta_1}$ for all $[S^*] \geq 0$, since $\delta_2 > \delta_1$. Thus $\max([S]_T) = \frac{Q}{\delta_1}$ when $[S^*] = 0$. Similarly, from eq. (16), $[S]_T \geq \frac{Q}{\delta_2}$ for all $[S] \geq 0$, since $\delta_2 > \delta_1$. Thus $\min([S]_T) = \frac{Q}{\delta_2}$ when $[S] = 0$.

We can plug in and solve for $\alpha = [S^*]/[S]_T$ when $\beta = [S]_T = \frac{1}{2}(\min [S]_T + \max [S]_T)$:

$$\frac{\min [S]_T + \max [S]_T}{2} = \frac{Q}{\delta_1} + \left(1 - \frac{\delta_2}{\delta_1}\right)[S^*] \implies \frac{Q}{2}\left(\frac{1}{\delta_1} + \frac{1}{\delta_2}\right) = \frac{Q}{\delta_1} + \left(1 - \frac{\delta_2}{\delta_1}\right)[S^*] \implies [S^*] = \frac{Q}{2\delta_2}$$

$$\alpha = \frac{[S^*]}{[S]_T} = \frac{Q/(2\delta_2)}{\frac{Q}{2}(1/\delta_1 + 1/\delta_2)} = \frac{\delta_1}{\delta_1 + \delta_2}$$

Substituting $\alpha = \dfrac{\delta_1}{\delta_1 + \delta_2}$ in eq. (13) and simplifying the resulting expression (3) yields:

$$r_{50} = \frac{Q + 2K_M\delta_1}{Q + 2K_M\delta_2}\left(1 + \frac{Q\delta_2}{k_{cat,D}[D]_T(\delta_2 + \delta_2)} + \frac{\delta_2(K_M + [D]_T)}{k_{cat,D}[D]_T}\right) \tag{17}$$

##### 1.9 Analytical expression for $n_{eff}$ of $[S]_T$ in Full model

To obtain an expression for $n_{eff}$, we need to first derive $EC_{10}$ and $EC_{90}$. Note that the 10% response for total substrate occurs at $\frac{9}{10}\min[S]_T + \frac{1}{10}\max[S]_T$ and the 90% response for total substrate occurs at $\frac{1}{10}\min[S]_T + \frac{9}{10}\max[S]_T$.

To derive $EC_{10}(\beta)$, we can plug in and solve for $\alpha$ when $\beta = \frac{9}{10}\min[S]_T + \frac{1}{10}\max[S]_T$:

$$\frac{9}{10}\min[S]_T + \frac{1}{10}\max[S]_T = \frac{Q}{\delta_1} + \left(1 - \frac{\delta_2}{\delta_1}\right)[S^*] \implies \frac{9Q}{10\delta_2} + \frac{Q}{10\delta_1} = \frac{Q}{\delta_1} + \left(1 - \frac{\delta_2}{\delta_1}\right)[S^*]$$
$$\implies [S^*] = \frac{9}{10}\left(\frac{Q}{\delta_2}\right)$$

$$\alpha = \frac{[S^*]}{[S]_T} = \frac{9Q/(10\delta_2)}{\frac{Q}{10}(1/\delta_1 + 9/\delta_2)} = \frac{9\delta_1}{9\delta_1 + \delta_2}$$

Substituting $\alpha = \dfrac{9\delta_1}{9\delta_1 + \delta_2}$ in eq. (13) and simplifying the resulting expression (3) yields:

$$EC_{10} = \frac{9(Q + 10K_M\delta_1)(9Q + 10[D]_T k_{cat,D} + 10K_M\delta_2 + 10[D]_T\delta_2)}{10[D]_T k_{cat,D}(9Q + 10K_M\delta_2)}$$

Similar calculations can be done to derive $EC_{90}$:

$$EC_{90} = \frac{(9Q + 10K_M\delta_1)(Q + 10[D]_T k_{cat,D} + 10K_M\delta_2 + 10[D]_T\delta_2)}{90[D]_T k_{cat,D}(Q + 10K_M\delta_2)}$$

Thus $n_{eff} = \log(81)/\log\left(\dfrac{EC_{90}}{EC_{10}}\right)$

$$= \frac{\log(81)}{\log(81) + \log\left(\frac{(Q + 10K_M\delta_1)(Q + 10K_M\delta_2)(10k_{cat,D}[D]_T + 9Q + 10(K_M + [D]_T)\delta_2)}{(9Q + 10K_M\delta_1)(9Q + 10K_M\delta_2)(10k_{cat,D}[D]_T + Q + 10(K_M + [D]_T)\delta_2)}\right)}$$

##### 1.9.1 Miscellaneous figures

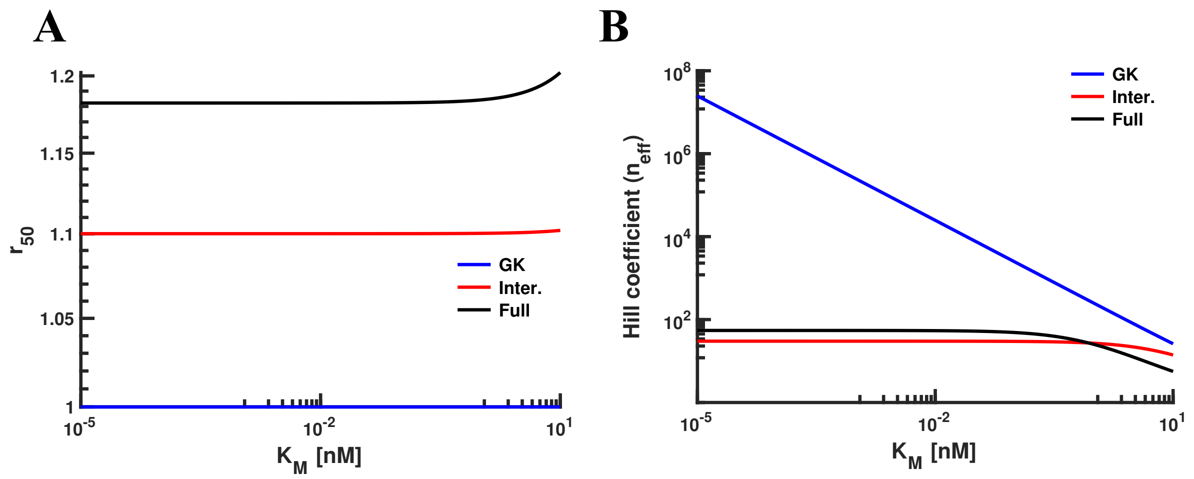

Figure S2: **Effects on the $r_{50}$ and $n_{eff}$ of various single-substrate models when $K_M$ is varied.** (A) Increasing $K_M$ has negligible effect on the $r_{50}$ for each of the models. $K_M$ axis in log scale. (B) Increasing $K_M$ results in a dramatic reduction of $n_{eff}$ for the GK model. For systems that incorporate protein turnover (i.e. the Intermediate and Full models), the reduction effect occurs for large enough $K_M$.

### 2 Single Substrate, Multiple Modification States

We generalize the model in the previous section to various models with arbitrarily long chains of ubiquitin units. In these models, substrates with 0 to 3 ubiquitin units attached are degraded at the first-order rate $\delta_1$ and substrates with more than 4 units are degraded at the rate $\delta_2 > \delta_1$. In what follows, the maximal length of the chain is denoted by $\ell$. The indices $i$ and $j$ represent the number of the ubiquitin unit and range from 0 to 3 and 4 to $\ell$ respectively. The index $x$ ranges from 1 to 3.

#### 2.1 Enzymatic reaction schemes

**Model with Distributive E3 & Trunk DUB**:

$$M + S^{(0)} \underset{k_{-,M}}{\overset{k_{+,M}}{\rightleftharpoons}} MS^{(0)} \xrightarrow{k_{cat,M}} M + S^{(1)}$$

$$M + S^{(1)} \underset{k_{-,M}}{\overset{k_{+,M}}{\rightleftharpoons}} MS^{(1)} \xrightarrow{k_{cat,M}} M + S^{(2)}$$

$$\vdots$$

$$M + S^{(\ell-1)} \underset{k_{-,M}}{\overset{k_{+,M}}{\rightleftharpoons}} MS^{(\ell-1)} \xrightarrow{k_{cat,M}} M + S^{(\ell)}$$

$$D + S^{(1)} \underset{k_{-,D}}{\overset{k_{+,D}}{\rightleftharpoons}} DS^{(1)} \xrightarrow{k_{cat,D}} D + S^{(0)}$$

$$D + S^{(2)} \underset{k_{-,D}}{\overset{k_{+,D}}{\rightleftharpoons}} DS^{(2)} \xrightarrow{k_{cat,D}} D + S^{(0)}$$

$$\vdots$$

$$D + S^{(\ell)} \underset{k_{-,D}}{\overset{k_{+,D}}{\rightleftharpoons}} DS^{(\ell)} \xrightarrow{k_{cat,D}} D + S^{(0)}$$

$$\emptyset \xrightarrow{Q} S^{(0)}$$

$$S^{(i)} \xrightarrow{\delta_1} \emptyset$$

$$S^{(j)} \xrightarrow{\delta_2} \emptyset$$

$$MS^{(i)} \xrightarrow{\delta_1} M$$

$$MS^{(j)} \xrightarrow{\delta_2} M$$

$$DS^{(x)} \xrightarrow{\delta_1} D$$

$$DS^{(j)} \xrightarrow{\delta_2} D$$

**Model with Distributive E3 & Sequential DUB**:

$$M + S^{(0)} \underset{k_{-,M}}{\overset{k_{+,M}}{\rightleftharpoons}} MS^{(0)} \xrightarrow{k_{cat,M}} M + S^{(1)}$$

$$M + S^{(1)} \underset{k_{-,M}}{\overset{k_{+,M}}{\rightleftharpoons}} MS^{(1)} \xrightarrow{k_{cat,M}} M + S^{(2)}$$

$$\vdots$$

$$M + S^{(\ell-1)} \underset{k_{-,M}}{\overset{k_{+,M}}{\rightleftharpoons}} MS^{(\ell-1)} \xrightarrow{k_{cat,M}} M + S^{(\ell)}$$

$$D + S^{(1)} \underset{k_{-,D}}{\overset{k_{+,D}}{\rightleftharpoons}} DS^{(1)} \xrightarrow{k_{cat,D}} D + S^{(0)}$$

$$D + S^{(2)} \underset{k_{-,D}}{\overset{k_{+,D}}{\rightleftharpoons}} DS^{(2)} \xrightarrow{k_{cat,D}} D + S^{(1)}$$

$$\vdots$$

$$D + S^{(\ell)} \underset{k_{-,D}}{\overset{k_{+,D}}{\rightleftharpoons}} DS^{(\ell)} \xrightarrow{k_{cat,D}} D + S^{(\ell-1)}$$

$$\emptyset \xrightarrow{Q} S^{(0)}$$

$$S^{(i)} \xrightarrow{\delta_1} \emptyset$$

$$S^{(j)} \xrightarrow{\delta_2} \emptyset$$

$$MS^{(i)} \xrightarrow{\delta_1} M$$

$$MS^{(j)} \xrightarrow{\delta_2} M$$

$$DS^{(x)} \xrightarrow{\delta_1} D$$

$$DS^{(j)} \xrightarrow{\delta_2} D$$

**Model with Distributive E3 & Processive DUB**:

$$M + S^{(0)} \underset{k_{-,M}}{\overset{k_{+,M}}{\rightleftharpoons}} MS^{(0)} \xrightarrow{k_{cat,M}} M + S^{(1)}$$

$$M + S^{(1)} \underset{k_{-,M}}{\overset{k_{+,M}}{\rightleftharpoons}} MS^{(1)} \xrightarrow{k_{cat,M}} M + S^{(2)}$$

$$\vdots$$

$$M + S^{(\ell-1)} \underset{k_{-,M}}{\overset{k_{+,M}}{\rightleftharpoons}} MS^{(\ell-1)} \xrightarrow{k_{cat,M}} M + S^{(\ell)}$$

$$D + S^{(1)} \underset{k_{-,D}}{\overset{k_{+,D}}{\rightleftharpoons}} DS^{(1)} \xrightarrow{k_{cat,D}} D + S^{(0)}$$

$$D + S^{(2)} \underset{k_{-,D}}{\overset{k_{+,D}}{\rightleftharpoons}} DS^{(2)} \xrightarrow{k_{cat,D}} DS^{(1)}$$

$$\vdots$$

$$D + S^{(\ell)} \underset{k_{-,D}}{\overset{k_{+,D}}{\rightleftharpoons}} DS^{(\ell)} \xrightarrow{k_{cat,D}} DS^{(\ell-1)}$$

$$\emptyset \xrightarrow{Q} S^{(0)}$$

$$S^{(i)} \xrightarrow{\delta_1} \emptyset$$

$$S^{(j)} \xrightarrow{\delta_2} \emptyset$$

$$MS^{(i)} \xrightarrow{\delta_1} M$$

$$MS^{(j)} \xrightarrow{\delta_2} M$$

$$DS^{(x)} \xrightarrow{\delta_1} D$$

$$DS^{(j)} \xrightarrow{\delta_2} D$$

**Model with Processive E3 & Trunk DUB**:

$$M + S^{(0)} \underset{k_{-,M}}{\overset{k_{+,M}}{\rightleftharpoons}} MS^{(0)} \xrightarrow{k_{cat,1,M}} MS^{(1)}$$

$$M + S^{(1)} \underset{k_{-,M}}{\overset{k_{+,M}}{\rightleftharpoons}} MS^{(1)} \xrightarrow{k_{cat,2,M}} MS^{(2)}$$

$$\vdots$$

$$M + S^{(\ell-1)} \underset{k_{-,M}}{\overset{k_{+,M}}{\rightleftharpoons}} MS^{(\ell-1)} \xrightarrow{k_{cat,2,M}} MS^{(\ell)}$$

$$D + S^{(1)} \underset{k_{-,D}}{\overset{k_{+,D}}{\rightleftharpoons}} DS^{(1)} \xrightarrow{k_{cat,D}} D + S^{(0)}$$

$$D + S^{(2)} \underset{k_{-,D}}{\overset{k_{+,D}}{\rightleftharpoons}} DS^{(2)} \xrightarrow{k_{cat,D}} D + S^{(0)}$$

$$\vdots$$

$$D + S^{(\ell)} \underset{k_{-,D}}{\overset{k_{+,D}}{\rightleftharpoons}} DS^{(\ell)} \xrightarrow{k_{cat,D}} D + S^{(0)}$$

$$\emptyset \xrightarrow{Q} S^{(0)}$$

$$S^{(i)} \xrightarrow{\delta_1} \emptyset$$

$$S^{(j)} \xrightarrow{\delta_2} \emptyset$$

$$MS^{(i)} \xrightarrow{\delta_1} M$$

$$MS^{(j)} \xrightarrow{\delta_2} M$$

$$DS^{(x)} \xrightarrow{\delta_1} D$$

$$DS^{(j)} \xrightarrow{\delta_2} D$$

**Model with Processive E3 & Sequential DUB**:

$$M + S^{(0)} \underset{k_{-,M}}{\overset{k_{+,M}}{\rightleftharpoons}} MS^{(0)} \xrightarrow{k_{cat,1,M}} MS^{(1)}$$

$$M + S^{(1)} \underset{k_{-,M}}{\overset{k_{+,M}}{\rightleftharpoons}} MS^{(1)} \xrightarrow{k_{cat,2,M}} MS^{(2)}$$

$$\vdots$$

$$M + S^{(\ell-1)} \underset{k_{-,M}}{\overset{k_{+,M}}{\rightleftharpoons}} MS^{(\ell-1)} \xrightarrow{k_{cat,2,M}} MS^{(\ell)}$$

$$D + S^{(1)} \underset{k_{-,D}}{\overset{k_{+,D}}{\rightleftharpoons}} DS^{(1)} \xrightarrow{k_{cat,D}} D + S^{(0)}$$

$$D + S^{(2)} \underset{k_{-,D}}{\overset{k_{+,D}}{\rightleftharpoons}} DS^{(2)} \xrightarrow{k_{cat,D}} D + S^{(1)}$$

$$\vdots$$

$$D + S^{(\ell)} \underset{k_{-,D}}{\overset{k_{+,D}}{\rightleftharpoons}} DS^{(\ell)} \xrightarrow{k_{cat,D}} D + S^{(\ell-1)}$$

$$\emptyset \xrightarrow{Q} S^{(0)}$$

$$S^{(i)} \xrightarrow{\delta_1} \emptyset$$

$$S^{(j)} \xrightarrow{\delta_2} \emptyset$$

$$MS^{(i)} \xrightarrow{\delta_1} M$$

$$MS^{(j)} \xrightarrow{\delta_2} M$$

$$DS^{(x)} \xrightarrow{\delta_1} D$$

$$DS^{(j)} \xrightarrow{\delta_2} D$$

**Model with Processive E3 & Processive DUB**:

$$M + S^{(0)} \underset{k_{-,M}}{\overset{k_{+,M}}{\rightleftharpoons}} MS^{(0)} \xrightarrow{k_{cat,1,M}} MS^{(1)}$$

$$M + S^{(1)} \underset{k_{-,M}}{\overset{k_{+,M}}{\rightleftharpoons}} MS^{(1)} \xrightarrow{k_{cat,2,M}} MS^{(2)}$$

$$\vdots$$

$$M + S^{(\ell-1)} \underset{k_{-,M}}{\overset{k_{+,M}}{\rightleftharpoons}} MS^{(\ell-1)} \xrightarrow{k_{cat,2,M}} MS^{(\ell)}$$

$$D + S^{(1)} \underset{k_{-,D}}{\overset{k_{+,D}}{\rightleftharpoons}} DS^{(1)} \xrightarrow{k_{cat,D}} D + S^{(0)}$$

$$D + S^{(2)} \underset{k_{-,D}}{\overset{k_{+,D}}{\rightleftharpoons}} DS^{(2)} \xrightarrow{k_{cat,D}} DS^{(1)}$$

$$\vdots$$

$$D + S^{(\ell)} \underset{k_{-,D}}{\overset{k_{+,D}}{\rightleftharpoons}} DS^{(\ell)} \xrightarrow{k_{cat,D}} DS^{(\ell-1)}$$

$$\emptyset \xrightarrow{Q} S^{(0)}$$

$$S^{(i)} \xrightarrow{\delta_1} \emptyset$$

$$S^{(j)} \xrightarrow{\delta_2} \emptyset$$

$$MS^{(i)} \xrightarrow{\delta_1} M$$

$$MS^{(j)} \xrightarrow{\delta_2} M$$

$$DS^{(x)} \xrightarrow{\delta_1} D$$

$$DS^{(j)} \xrightarrow{\delta_2} D$$

##### 2.2 Stochastic simulations

Note that the definition of a maximum length $\ell$ above is necessary in order to ensure a finite set of chemical reactions and ODEs. To determine if this truncation has any effect on the results, we compared our deterministic case to stochastic simulations in which we allow the ubiquitin chains to reach an arbitrary length. Our approach to developing these simulations is inspired by "agent-based" simulators developed for the stochastic simulation of rule-based models (6; 7). Due to technical considerations with the mechanism, however, we wrote our own dedicated code for these simulations, following closely the approach taken in our previous work on modeling length control in the bacterial Type III Secretion System (8).

Briefly, our simulations contain three types of agents: the M enzyme, the D enzyme, and the substrate S. These agents are represented *independently*; in other words, if there are 1000 S molecules in the simulation, this is represented by having 1000 distinct "S" agents in memory. Each S agent has associated with it a number that represents the length of its ubiquitin chain. These lengths can range from 0 to the largest number that can be represented by the particular data structure. Since this number is much, much larger than the largest value ever practically observed in the simulation, this essentially corresponds to allowing for arbitrary chain lengths.

We wrote a separate simulation in C++ for all of the scenarios described above. All of the parameters from the deterministic simulations were converted to their corresponding stochastic values in a straightforward way (9). Since these simulations are relatively expensive, we performed simulations for a subset of the parameters considered in our deterministic simulations (see below). All simulation codes are available upon request.

##### 2.3 Graphical results

In the following figures, in order to choose a reasonable value for $\ell$, we first ran simulations such that changes in the $r_{50}$ and $n_{eff}$ were negligible beyond a point. Using these results, we then chose $\ell = 500$ by inspection.

The red curves correspond to the case of saturation, while the blue curves correspond to the unsaturated case. The lines indicate the numerically integrated deterministic solutions and the dots indicate averages from agent-based stochastic simulations.

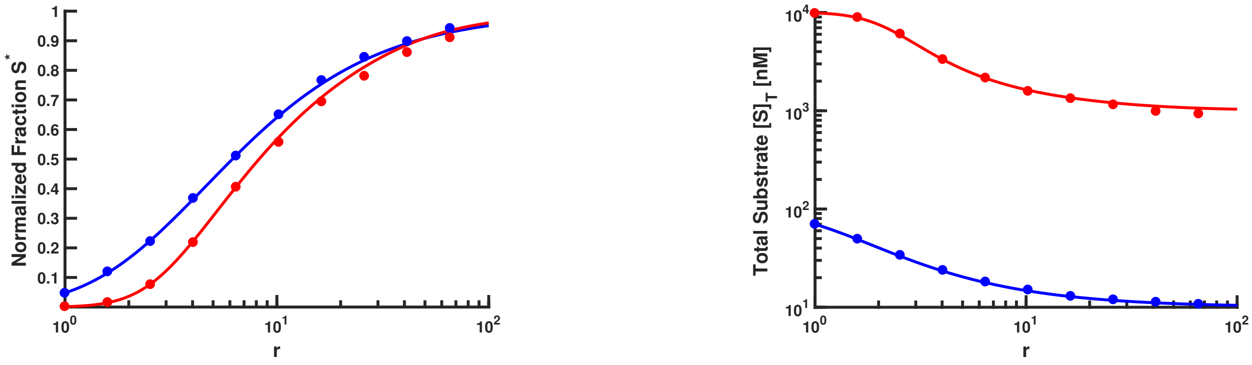

Figure S3: **Model with Distributive E3 & Trunk DUB.**

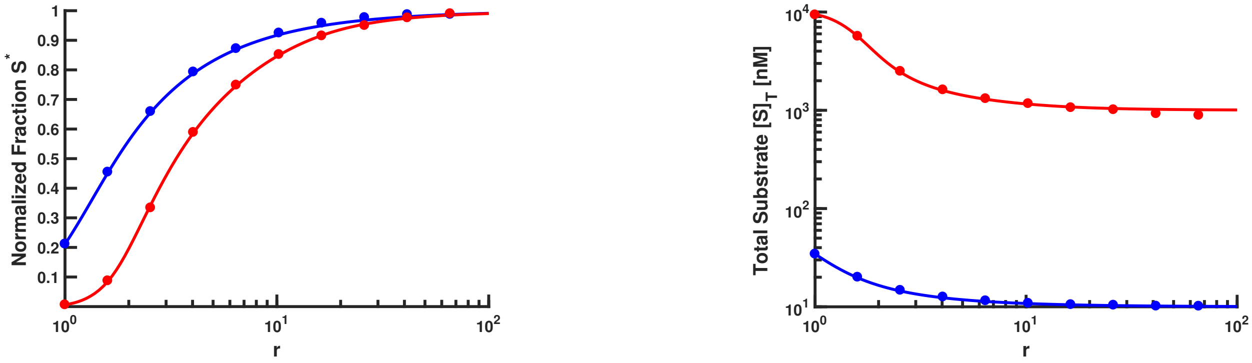

Figure S4: **Model with Distributive E3 & Sequential DUB.**

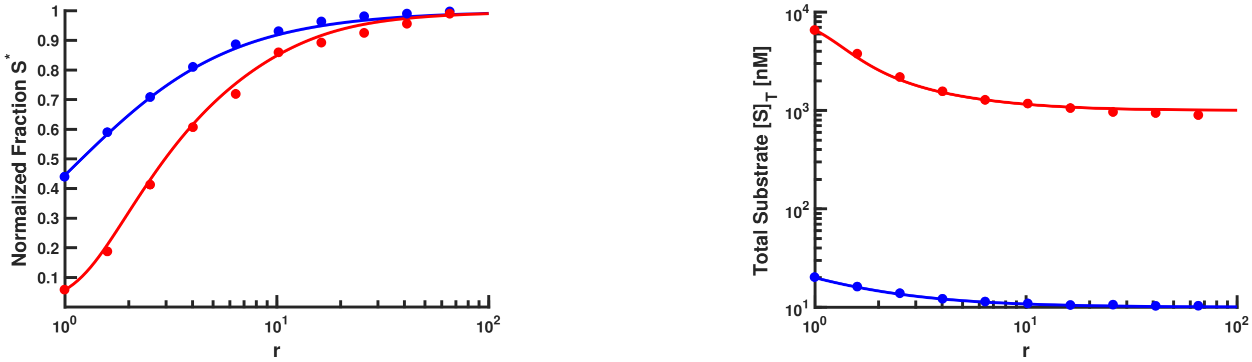

Figure S5: **Model with Distributive E3 & Processive DUB.**

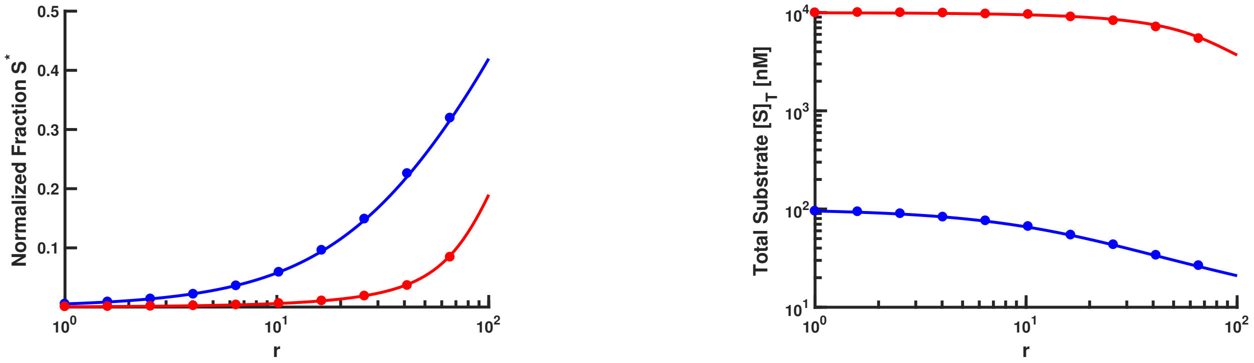

Figure S6: **Model with Processive E3 & Trunk DUB.**

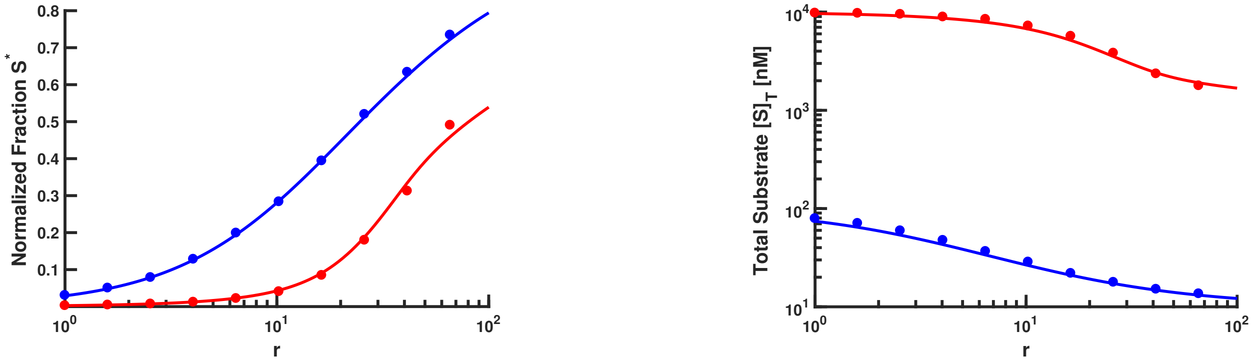

Figure S7: **Model with Processive E3 & Processive DUB.**

#### 3 Multiple Substrates, Single Modification State

The subscript index $z$ represents the substrate number, ranging from 1 to $N$, where $N$ is the total number of substrates.

$$M + S_z \underset{k_{-,M}}{\overset{k_{+,M}}{\rightleftharpoons}} MS_z \xrightarrow{k_{cat,M}} M + S_z^*$$

$$D + S_z^* \underset{k_{-,D}}{\overset{k_{+,D}}{\rightleftharpoons}} DS_z^* \xrightarrow{k_{cat,D}} D + S_z$$

$$\emptyset \xrightarrow{Q_z} S$$

$$S_z \xrightarrow{\delta_1} \emptyset$$

$$S_z^* \xrightarrow{\delta_2} \emptyset$$

$$MS_z \xrightarrow{\delta_1} M$$

$$DS_z^* \xrightarrow{\delta_2} D$$

#### 3.1 Equations

##### 3.1.1 Main

$$\frac{d[S_z]}{dt} = Q_z - k_{+,M}[M][S_z] + k_{-,M}[MS_z] + k_{cat,D}[DS_z^*] - \delta_1[S_z] \tag{18a}$$

$$\frac{d[S_z^*]}{dt} = -k_{+,D}[D][S_z^*] + k_{-,D}[DS_z^*] + k_{cat,M}[MS_z] - \delta_2[S_z^*] \tag{18b}$$

$$\frac{d[MS_z]}{dt} = k_{+,M}[M][S_z] - (k_{-,M} + k_{cat,M} + \delta_1)[MS_z] \tag{18c}$$

$$\frac{d[DS_z^*]}{dt} = k_{+,D}[D][S_z^*] - (k_{-,D} + k_{cat,D} + \delta_2)[DS_z^*] \tag{18d}$$

$$\frac{d[M]}{dt} = (k_{-,M} + k_{cat,M} + \delta_1)\sum_{z=1}^{N}[MS_z] - k_{+,M}[M]\sum_{z=1}^{N}[S_z] \tag{18e}$$

$$\frac{d[D]}{dt} = (k_{-,D} + k_{cat,D} + \delta_2)\sum_{z=1}^{N}[DS_z^*] - k_{+,D}[D]\sum_{z=1}^{N}[S_z^*] \tag{18f}$$

##### 3.1.2 Mass conservation

$$[M]_T = [M] + \sum_{z=1}^{N}[MS_z]$$

$$[D]_T = [D] + \sum_{z=1}^{N}[DS_z^*]$$

$$[S]_T = \sum_{z=1}^{N}([S_z] + [S_z^*] + [MS_z] + [DS_z^*])$$

#### 3.2 Comments

Adding multiple substrates to the system results in additive effects. Similar conclusions hold for multiple substrates with just one modification state as for one substrate with one modification state (Section 1). The only changes in the relevant expressions are:

(i) The substrate production rate $Q$ becomes $\sum_z Q_z$

(ii) $S$ becomes $\sum_z S_z$

(iii) $S^*$ becomes $\sum_z S_z^*$

##### 3.3 Analytical expression for $r_{50}([S_1]_T)$

By inspecting eq. (13) and its derivation, one can see that the expression for $r$ in this case will be similar. Following the derivation in Section 1.8, we can obtain $r_{50}([S_1]_T)$ as follows:

$$r_{50}([S_1]_T) = \frac{\sum_{i=1}^{N} \frac{\alpha_i Q_i}{\delta_1 + \alpha_i(\delta_2 - \delta_1)}}{k_{cat,D}} \left( \frac{\delta_2}{[D]_T} + \frac{\delta_2 + k_{cat,D}}{K_M + \sum_{i=1}^{N} \frac{\alpha_i Q_i}{\delta_1 + \alpha_i(\delta_2 - \delta_1)}} \right) \left( 1 + \frac{K_M}{\sum_{i=1}^{N} \frac{(1-\alpha_i) Q_i}{\delta_1 + \alpha_i(\delta_2 - \delta_1)}} \right)$$

where

$$\alpha_1 = \frac{\delta_1}{\delta_1 + \delta_2}$$

$$\alpha_i = \frac{[S_i^*]}{[S_i]_T} \text{ corresponding to } \alpha_1$$

$$N = \text{Total Number of Substrates}$$

For the sake of illustration, the figure below is for $N = 10$. The semi-analytical curve was obtained by substituting values of $\alpha_i$ obtained empirically from simulation into the analytical expression for $r_{50}([S_1]_T)$.

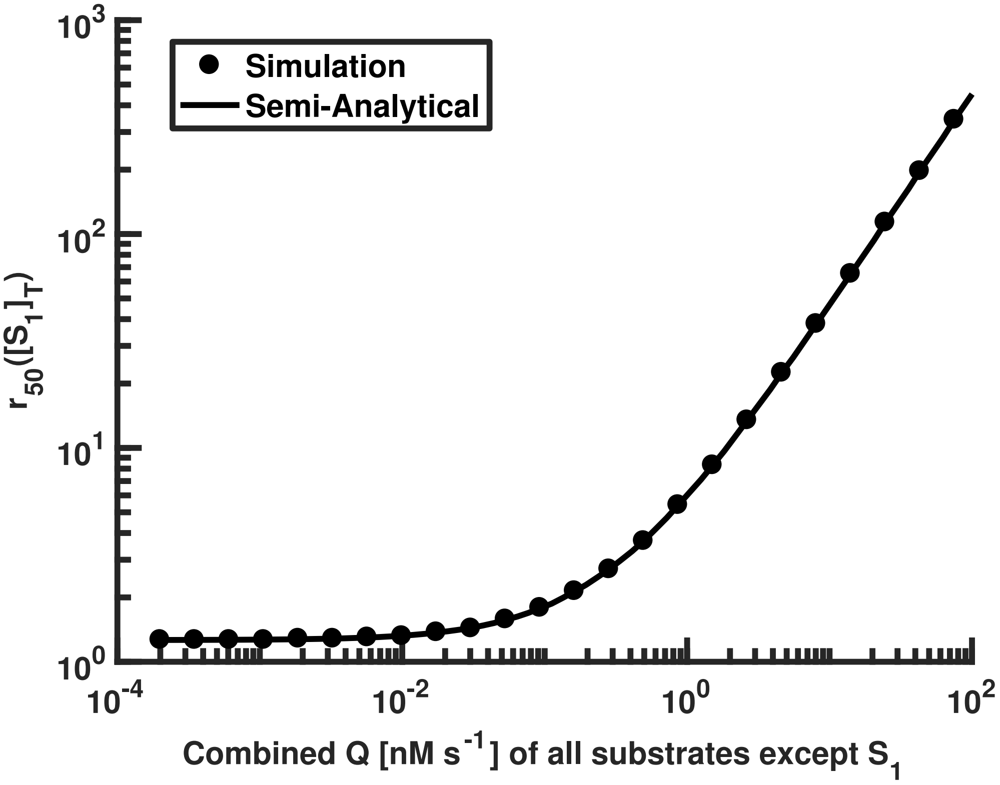

#### 4 Multiple Substrates, Multiple Modification States

We generalize the model in Section 2 to a model with multiple substrates. For the sake of illustration, the model with Processive E3 & Sequential DUB is described by the following schematic of enzymatic reactions:

**Model with Processive E3 & Sequential DUB**:

$$M + S_z^{(0)} \underset{k_{-,M}}{\overset{k_{+,M}}{\rightleftharpoons}} MS_z^{(0)} \xrightarrow{k_{cat,M}} MS_z^{(1)}$$

$$M + S_z^{(1)} \underset{k_{-,M}}{\overset{k_{+,M}}{\rightleftharpoons}} MS_z^{(1)} \xrightarrow{k_{cat,M}} MS_z^{(2)}$$

$$\vdots$$

$$M + S_z^{(\ell-1)} \underset{k_{-,M}}{\overset{k_{+,M}}{\rightleftharpoons}} MS_z^{(\ell-1)} \xrightarrow{k_{cat,M}} MS_z^{(\ell)}$$

$$D + S_z^{(1)} \underset{k_{-,D}}{\overset{k_{+,D}}{\rightleftharpoons}} DS_z^{(1)} \xrightarrow{k_{cat,D}} D + S_z^{(0)}$$

$$D + S_z^{(2)} \underset{k_{-,D}}{\overset{k_{+,D}}{\rightleftharpoons}} DS_z^{(2)} \xrightarrow{k_{cat,D}} D + S_z^{(1)}$$

$$\vdots$$

$$D + S_z^{(\ell)} \underset{k_{-,D}}{\overset{k_{+,D}}{\rightleftharpoons}} DS_z^{(\ell)} \xrightarrow{k_{cat,D}} D + S_z^{(\ell-1)}$$

$$\emptyset \xrightarrow{Q} S_z^{(0)}$$

$$S_z^{(i)} \xrightarrow{\delta_1} \emptyset$$

$$S_z^{(j)} \xrightarrow{\delta_2} \emptyset$$

$$MS_z^{(i)} \xrightarrow{\delta_1} M$$

$$MS_z^{(j)} \xrightarrow{\delta_2} M$$

$$DS_z^{(x)} \xrightarrow{\delta_1} D$$

$$DS_z^{(j)} \xrightarrow{\delta_2} D$$

2. L. A. Segel, L. Edelstein-Keshet, *A Primer on Mathematical Models in Biology* (Society for

Industrial and Applied Mathematics, Philadelphia, PA, 2013).

3. Wolfram Research, Inc., Mathematica 10.0.

4. W. Dubitzky, O. Wolkenhauer, H. Yokota, K.-H. Cho, *Encyclopedia of Systems Biology* (Springer, New York, NY, 2013).
